# Transcriptomic Effects of Low-Dose Inorganic Arsenic Exposure on Murine Bone Marrow-Derived Macrophages

**DOI:** 10.1101/2023.07.26.550543

**Authors:** Emily J. Illingworth, Alexandra Maertens, Fenna C.M. Sillé

**Affiliations:** Department of Environmental Health and Engineering, Bloomberg School of Public Health, Johns Hopkins University, Baltimore, MD, USA

**Keywords:** Arsenic, Immunotoxicology, Macrophages, Transcriptome, Sex Differences

## Abstract

Both tissue-resident macrophages and monocytes recruited from the bone marrow that transform into tissue-resident cells play critical roles in mediating homeostasis as well as in the pathology of inflammatory diseases. Inorganic arsenic (iAs) is the most common drinking water contaminant worldwide and represents a major public health concern. Several diseases that macrophages have implicated involvement in are caused by iAs exposure, including cardiovascular disease, cancer, and increased risk of infectious disease. Therefore, understanding the effects of iAs exposure on macrophages can help us better grasp the full range of arsenic immunotoxicity and better design therapeutic targets for iAs-induced diseases particularly in exposed populations. In this study, we analyzed the transcriptome of low dose iAs-exposed male and female murine bone marrow-derived macrophages (BMDMs) with either M0, M1, or M2 stimulation. We identified differentially expressed genes by iAs in a sex- and stimulation-dependent manner and used bioinformatics tools to predict protein-protein interactions, transcriptional regulatory networks, and associated biological processes. Overall, our data suggest that M1-stimulated, especially female-derived, BMDMs are most susceptible to iAs exposure. Most notably, we observed significant downregulation of major proinflammatory transcription factors, like IRF8, and its downstream targets, as well as genes encoding proteins involved in pattern recognition and antigen presentation, such as TLR7, TLR8, and H2-D1, potentially providing causal insight regarding arsenic’s role in perturbing immune responses to infectious diseases. We also observed significant downregulation of genes involved in processes crucial to coordinating a proinflammatory response including leukocyte migration, differentiation, and cytokine and chemokine production and response. Finally, we discovered that 24 X-linked genes were dysregulated in iAs-exposed female stimulation groups compared to only 3 across the iAs-exposed male stimulation groups. These findings elucidate the potential mechanisms underlying the sex-differential iAs-associated immune-related disease risk.

## INTRODUCTION

Inorganic arsenic (iAs) is one of the most common water contaminants worldwide. It is naturally occurring in groundwater and can therefore contaminate water used for drinking and crop irrigation. Globally, over 200 million people are exposed to iAs at levels above the World Health Organization safety standard of 10 µg/L (10 ppb) (Podgorski 2020; WHO 2006). In the US alone, 4.1 million people are estimated to be exposed to dangerous levels of iAs, a number that has almost doubled since 2017 in part due to drought conditions exacerbated by climate change (Lombard 2021). These exposures pose a major public health concern, as arsenic is an IARC-classified Group 1 carcinogen of the lung, skin, and bladder (IARC 2012), and causes increased risk for cardiovascular disease (Moon 2012; Yuan 2007) as well as adverse endocrinological (Hong-Jie 2016) and immune effects (Dangleben 2013).

Antigen presenting cells represent a group of immune cell targets of arsenic toxicity. Macrophages are professional antigen presenting cells that play important roles in linking the innate and adaptive immune system (Harding 2010; Zhang 2022). Tissue resident macrophages, along with other cells in the innate immune system such as dendritic cells, natural killer cells, and neutrophils, recognize local pathogen-associated molecular patterns (PAMPs) or damage-associated molecular patterns (DAMPs), and as such, represent the first line of defense against insult. Based on the type of stimuli they receive, these innate cells can phagocytose pathogens or tumor cells, secrete humoral factors, and help to orchestrate an appropriate response by recruiting other innate as well as adaptive immune cells to respond to the insult (Hajishengallis 2015; Hirayama 2018; Parameswaran 2010; Zhou 2021). For example, upon engagement of pattern recognition receptor (PRR) by viral pattern associated molecular patterns (PAMPs), such as Toll-like Receptor 7 (TLR7) with virus-derived ssRNA, specific transcription factors (e.g., Interferon regulatory factors [IRFs]) are induced that in turn control the transcription and translation of antiviral genes. In addition to clearing the virus, macrophages release pro inflammatory cytokines to recruit other phagocytes and effector T cells (Duan 2017). Apoptotic cells are also be recognized by PRRs on macrophages (Lauber 2012), which then clear cellular debris, and secrete anti-inflammatory cytokines such as TGFβ, IL-10, and arachidonic acid metabolites to help execute a “non-disruptive” cell death (Fogarty 2015; Gordon 2018).

Macrophages are highly plastic and exist on a spectrum of activation phenotypes depending on cues they receive from their microenvironment, with “M1” macrophages representing one end of the spectrum and “M2” macrophages representing the opposite (Mills 2000; Orecchinio 2019). M1 macrophages are pro-inflammatory and are classically polarized with lipopolysaccharide (LPS) plus interferon-gamma (IFNγ), while M2 macrophages resolve inflammation and are polarized by cytokines such as interleukin-4 (IL-4) plus interleukin-13 (IL-13). M1 and M2 macrophages are functionally and phenotypically distinct, as M1 macrophages express iNOS, CD80, and CD68 while M2 macrophages express CD163, Arg1, and each secrete distinct chemokine and cytokines (Mills 2000; Röszer 2015). Though the M1-M2 dichotomy is useful for discussion and experimental purposes, it should be noted that this dichotomy is extremely oversimplified, as macrophages exist as heterogenous phenotypes and can re-polarize upon interacting with new microenvironmental stimuli (Orecchioni 2019).

Bone marrow-derived macrophages (BMDMs) are a useful *in vitro* model for studying macrophage responses to stimuli (Zajd 2020). BMDMs are typically differentiated from bone marrow precursor cells in the presence of macrophage colony stimulating factor (M-CSF). *In vivo*, M-CSF is produced by endothelial cells, fibroblasts, and other macrophages in response to PAMPs and/or DAMPs (Ushach 2016). Monocytes produced in the bone marrow traffic through the blood to the sites of insult where they differentiate into mature macrophages based on further cues they receive in that microenvironment (Mills 2000; Orecchioni 2019). Monocytes act as one of the first responders during an inflammatory event; therefore, they are involved in a myriad of different diseases (Cho 2012; Davies 2013; Iwamoto 2021; Zajd 2020). Importantly, many diseases are associated with arsenic exposure including cancer (IARC 2012), atherosclerosis (Wang 2002), and increased susceptibility to infectious diseases (Liao 2011; Rahman 2011; Smith 2011).

Understanding the distinct effects of arsenic on monocytes and macrophages can help us pinpoint their roles in arsenic-induced diseases and may ultimately help us identify immunotherapeutic targets for prevention and treatment of these diseases in arsenic-exposed populations. In this study, we explored the effects of a noncytotoxic, and human-relevant concentration of iAs (0.1 µM sodium (meta) arsenite, which equates to approximately 13 ppb) on gene expression profiles of BMDMs from male and female mice in the context of M0, M1, and M2 stimulation using RNA-sequencing. We demonstrate that the effects of arsenic on BMDMs are sex- and stimulation-dependent and have distinct consequences on biological processes and gene regulatory networks related to immune function that may be relevant for developing interventions for arsenic-induced diseases. This work will inform hypothesis-driven research to tease apart the mechanisms by which arsenic affects macrophages in different disease contexts.

## METHODS

### Animals

Adult (8–10-week-old) male and female C57BL/6CR mice, purchased from Charles River Laboratories (Frederick, MD, USA), were given 1 week to acclimate before experiments. Mice were cohoused five/cage in a temperature- and humidity-controlled facility with a light:dark cycle maintained at 14:10 and provided food and drinking water *ad libitum*. Mice received low arsenic chow (AIN-93M: Research Diets, New Brunswick, NJ, CAT# D10012Mi) and arsenic-free water (Crystal Geiser, Calistoga, CA, USA). All animal procedures were conducted under Johns Hopkins University Animal Care and Use Committee protocol MO20H281.

### Isolation, Differentiation, and Polarization of Bone Marrow-derived Macrophages

Bone marrow from C57BL/6CR mice was isolated from the femur and tibia of 8–10-week-old mice, with bone marrow from male and female mice processed separately to maintain sex stratification for this study. Bones were flushed with PBS +10% FBS using a 23G needle and a single cell suspension was created by passing the cells through the needle and a 70 µm cell strainer. Red blood cells were lysed with ACK lysis buffer (Quality Biological, Gaithersburg, MD, USA) and pelleted cells were resuspended and plated with Dulbecco’s Modified Eagle Medium (DMEM) media containing 10% FBS, 200 µM L-glutamine, 100 µM HEPES (Gibco, Waltham, MA, USA), 100 µM MEM Non-Essential Amino Acids (NEAA) (Gibco, Waltham, MA, USA), 100 µM sodium pyruvate (Gibco, Waltham, MA, USA), and 25 ng/mL of macrophage colony stimulating factor (M-CSF) (complete media) as previously described (Assouvie 2018). Cells were not pooled across biological replicates. During the 7-day differentiation period, half the cells (∼2-4 x 10^7^) from each animal were maintained in medium supplemented with inorganic arsenic (iAs) in the form of 0.1 µM (∼13 ppb) sodium (meta) arsenite (Millipore Sigma, St. Louis, MO, USA, CAT# S7400-100G). Cultures were maintained at 37 °C with 5% CO_2_ for 7 days with medium refreshment (half of the media was replaced with 2x growth factors) +/-0.1 µM sodium (meta) arsenite after 4 days. After 7 days of culture, complete and arsenic-containing media was removed and adherent cells were incubated with ice-cold PBS and harvested by manual cell scraping, plated at 2-5 x 10^6^ cells per well in a clear U-bottom 96-well plate, then polarized overnight to the M0 (complete medium alone), the M1 (complete media containing 300 ng/mL LPS and 6.25 ng/mL IFNγ), or the M2 (complete media containing 20 ng/mL each of IL-4 and IL-13) phenotype. The cells were not exposed to sodium (meta) arsenite during polarization.

### RNA Sequencing

Total RNA was isolated from polarized murine bone marrow-derived macrophages using the Quick-RNA™ MiniPrep Kit (Zymo Research, Irvine, CA, USA). RNA quantity and quality was verified by two methods: using a NanoDrop™ 2000 spectrophotometer (Thermo Scientific™, Waltham, MA, USA) and with the Ribogreen assay (Quant-it™ RiboGreen RNA Assay Kit, Invitrogen™, Waltham, MA, USA). Preparation of RNA libraries and transcriptome sequencing was obtained using the Illumina NovoSeq platforms by Novogene Co (Sacramento, CA, USA). Briefly, samples were subjected to quality control before library construction with Nanodrop and agarose gel electrophoresis for potential RNA degradation or contamination. Quality control information for each sample, including RNA integrity number, is provided in **Table 2-1**. After paired-end sequencing, mRNA libraries were prepared with poly A enrichment using the NovaSeq 6000 PE150 platform by Novogene Co (Sacramento, CA, USA) with a minimum requirement of 6 G of raw data per sample. Raw data was filtered by removing reads containing adapters, removing reads containing >10% unresolvable bases and removing reads with >50% low quality bases. Reads were mapped to the reference *Mus musculus* genome. As proof of concept that we established 3 distinct stimulations using M0, M1, or M2 conditions, a principal component analysis plot showing all 48 samples sent for sequencing is provided (**Fig. 2-1**).

**Table 2-1.**
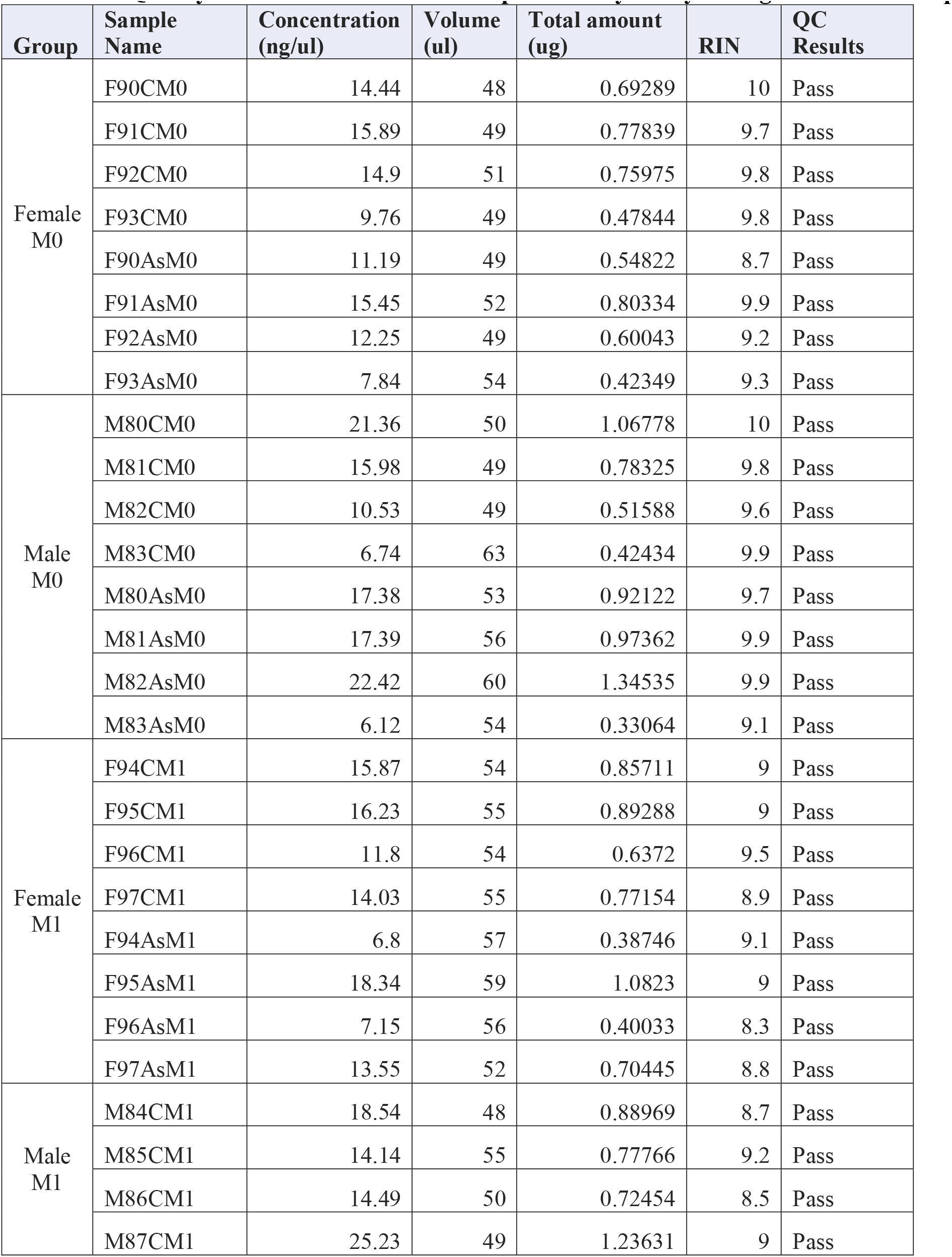
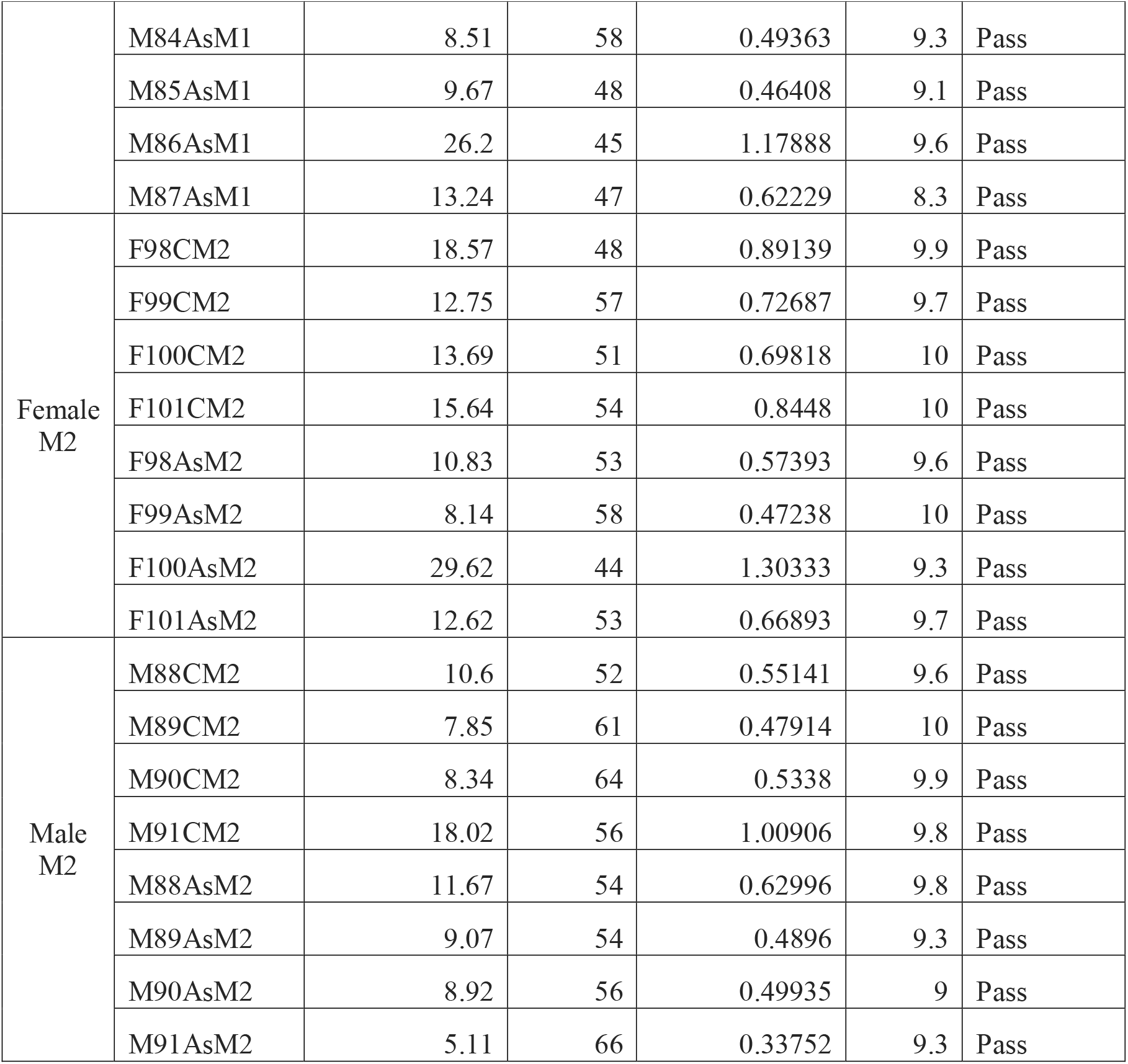
Quality control results of each sample as analyzed by Novogene for RNA-seq.

**Figure 2-1.**
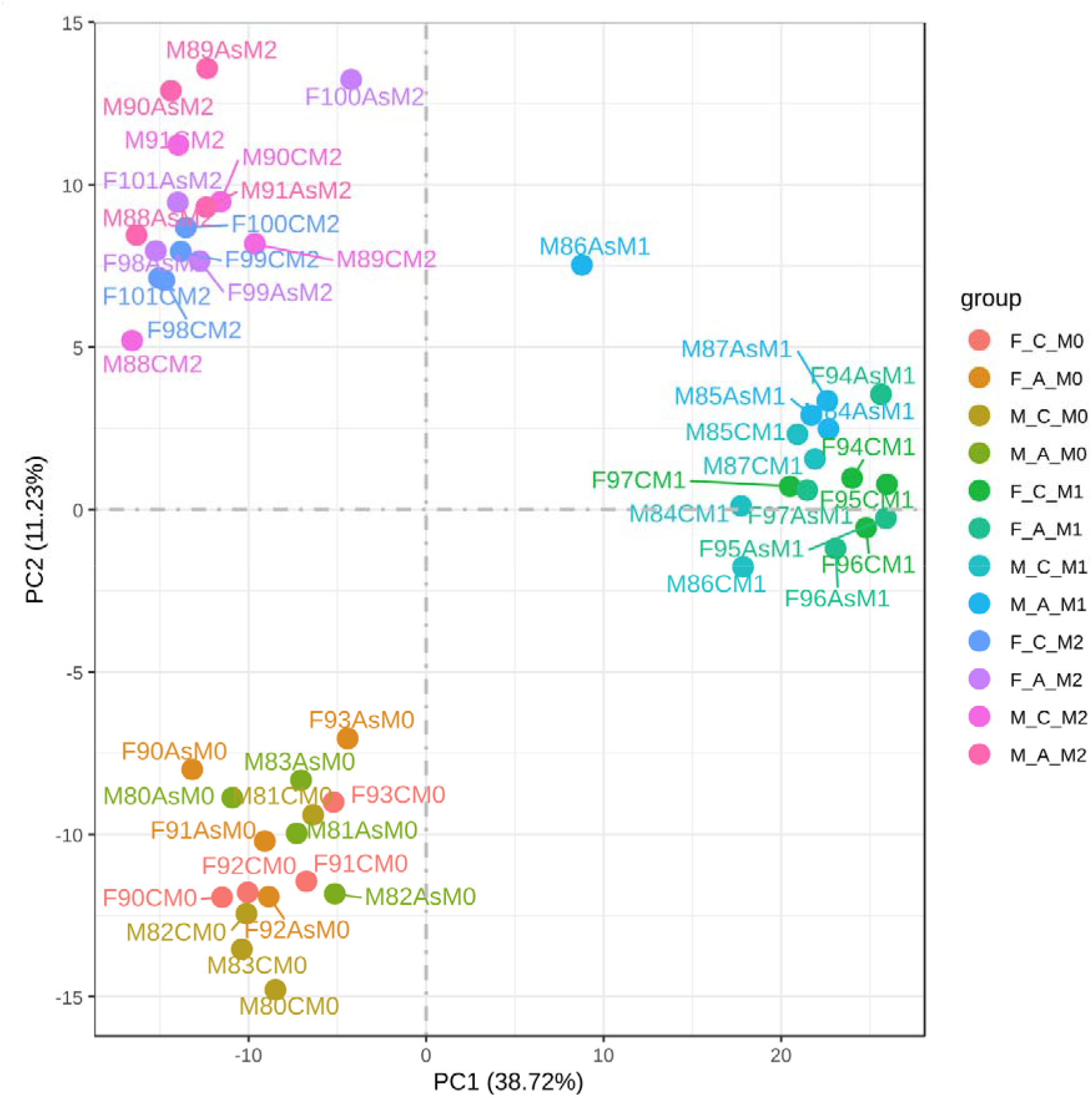
Principal component analysis (PCA) plots of M0, M1, and M2-stimulated BMDMs across both sexes for each sample sent for RNA-sequencing. Distinct clusters of each sample stimulation are shown by color coding across sexes and treatment. Sample name is given next to each plotted dot.

### Statistical analysis

RNA-seq reads were quantified and analyzed by Novogene (Novogene Co, Sacramento, CA, USA) using the R package DESeq2, which uses negative binomial generalized linear models to test for differential gene expression of biological replicates across sex and stimulation groups. The resulting *p-*values were adjusted using the Benjamini Hochberg approach in order to control the false discovery rate.

### STRING Database for Predicting Protein-protein Interaction Networks

Search Tool for the Retrieval of Interacting Genes/Proteins (STRING) is a database that extracts experimental and curated data from several sources to deduce protein-protein interaction (PPI) networks from a set of protein-coding genes (Szklarczyk 2021). Using our DEG list cutoff, |log2FC|>0 and adjusted *p-*value cutoff of <0.05, we generated PPI networks for the gene sets for each stimulation group across sexes using the STRING medium confidence required interaction score. We limited our network generation to only include experimental evidence and evidence from curated databases as active interaction sources. For visualization purposes, we chose to hide disconnected nodes in the network display settings. However, these disconnected nodes are considered in the full PPI list results.

### Visualizing STRING Maps in Cytoscape

Cytoscape (v 3.9.1) (Shannon 2003) an open-source platform, was used for general network visualization of STRING-generated protein-protein interactions. STRING network interaction tables were downloaded in tsv format and uploaded to Cytoscape. Source and target nodes were defined with edge attributes defined only by evidence scores from experimentally validated interactions and evidence scores from curated databases. Nodes with gene interactions that we identified as immunologically relevant were highlighted based on functional categories: integrin signaling (green), coagulation factors (red), chemotaxis and chemokine signaling (orange), pattern recognition and antigen presentation (pink), splicing machinery (yellow), lipid metabolism (purple), complement system (blue), and cytokine signaling (aqua). All Cytoscape maps are displayed using the degree sorted circle layout where the highest degree nodes (high connectivity “hubs”) are in the same region of the circle (lower right quadrant) and the degree decreases as you proceed counterclockwise around the circle.

### Fantom Database for Predicting Transcriptional Regulatory Networks

To generate transcriptional regulatory networks, we used Fantom EdgeExpressDB (Ravasi 2010; Severin 2009) subnet view. In the input settings, we removed generation of interactions by predicted transcription factor binding sites and miRNA predicted targets and visualized only primary edge types informed by published protein-DNA interactions, protein-protein interactions, chromatin immunoprecipitation data, and data from siRNA/miRNA perturbation experiments. For visualization purposes, singleton nodes were removed from the network.

### Gene Ontology Enrichment Analysis of Biological Processes, Cellular Components, and Molecular Function

Gene ontology (GO) enrichment analysis (Ashburner 2000; GO Consortium 2021; Mi 2019) for biological processes, cellular components, and molecular function was performed on each differentially expressed gene set for male and female mice. For inputting gene lists, we used a threshold of |log2FC|>0 and adjusted *p-*value cutoff of <0.05 for both up- and downregulated gene sets (relative to the iAs treatment group) across M0, M1, and M2 stimulation groups (for a total of 12 analyses per enrichment). The top 10 significantly GO enrichments from each analysis (or less if fewer than 10 endpoints were significant) were determined by Fischer’s Exact test with False Discovery Rate (FDR) for multiple hypothesis testing *p*-value adjustment. In addition, when there were more than 10 results, we looked at the complete list of statistically significant GO enriched biological processes and plotted additional hits based on immune relevance. If no statistically significant enrichment was found, nothing was plotted for that comparison.

## RESULTS

### iAs-induced Sex-specific Differential Gene Expression in BMDMs Across M0, M1, and M2 Stimulation Groups

iAs exposure during BMDM differentiation resulted in transcriptomic changes in both male- and female-derived cells across the M0, M1, and M2 stimulation groups. Using a cutoff of |log2FC| >0 and adjusted *p*-value of <0.05 to define “differential expression,” we identified a total of 1,543 differentially expressed genes (DEGs) altered in the Female M0 group (916 upregulated; 627 downregulated) (**Fig. 2-3A**), a total of 2,636 genes altered in the Female M1 group (1,354 upregulated; 1,282 downregulated) (**Fig. 2-2B**), and a total of 1,489 genes altered in the Female M2 group (819 upregulated; 670 downregulated) (**Fig. 2-2C**). In male BMDMs, a total of 1,195 genes altered in the Male M0 group were identified (768 upregulated; 427 downregulated) (**Fig. 2-2D**), a total of 1,573 genes altered in the Male M1 group (855 upregulated; 718 downregulated) (**Fig. 2-2E**), and a total of 951 genes altered in the Male M2 group (631 upregulated; 320 downregulated) (**Fig. 2-2F**). When we compared DEGs across stimulation groups for both sexes, we also found many shared genes across all stimulation groups (251 genes for females, and 45 genes for males), as well as those that are either shared between only two stimulation groups or uniquely altered in a single stimulation condition (**Fig. 2-3**). To highlight the top hits for each comparison group, the top 50 genes meeting more a stringent cutoff of |log2FC| >0.2, along with their gene descriptions from the Mouse Genome Informatics (MGI) database (Bult 2019), the adjusted *p-*values, and the chromosome locations are provided for male- and female-derived BMDMs (**Tables 2-1** − **2-12**).

**Figure 2-2.**
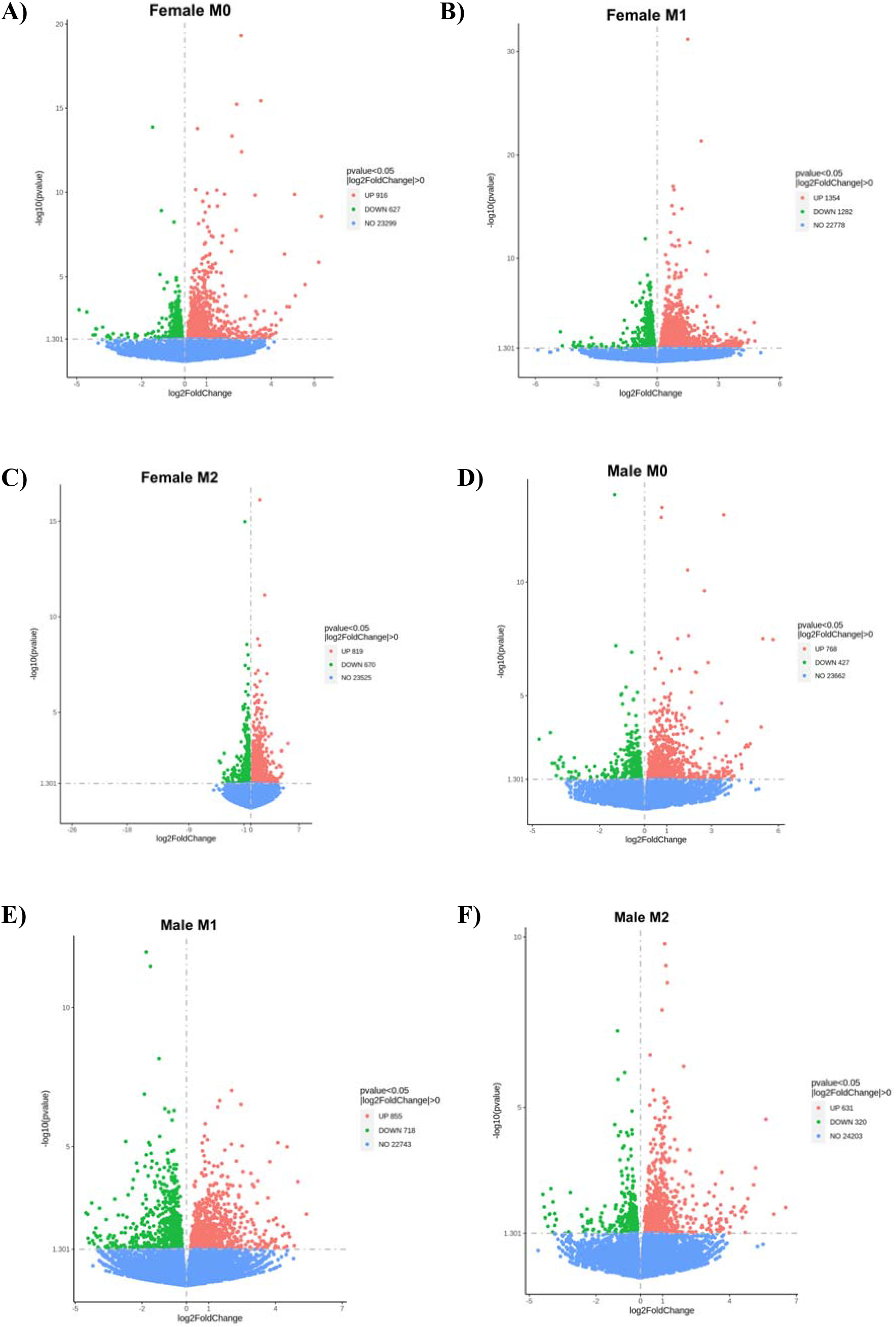
Volcano plots showing significantly up- or down-regulated genexposed female and male BMDMs. Volcano plots generated with our DEG cut-off |log2FC|>0 and adjusted p-value of <0.05 for Female M0 (A), Female M1 (B), Female M2 (C), Male M0 (D), Male M1 (E), and Male M2 (F) groups. Genes meeting this cut-off are represented by pink (upregulated genes) or green (downregulated genes) dots. Genes that did not meet this cut-off are shown as blue dots.

**Figure 2-3.**
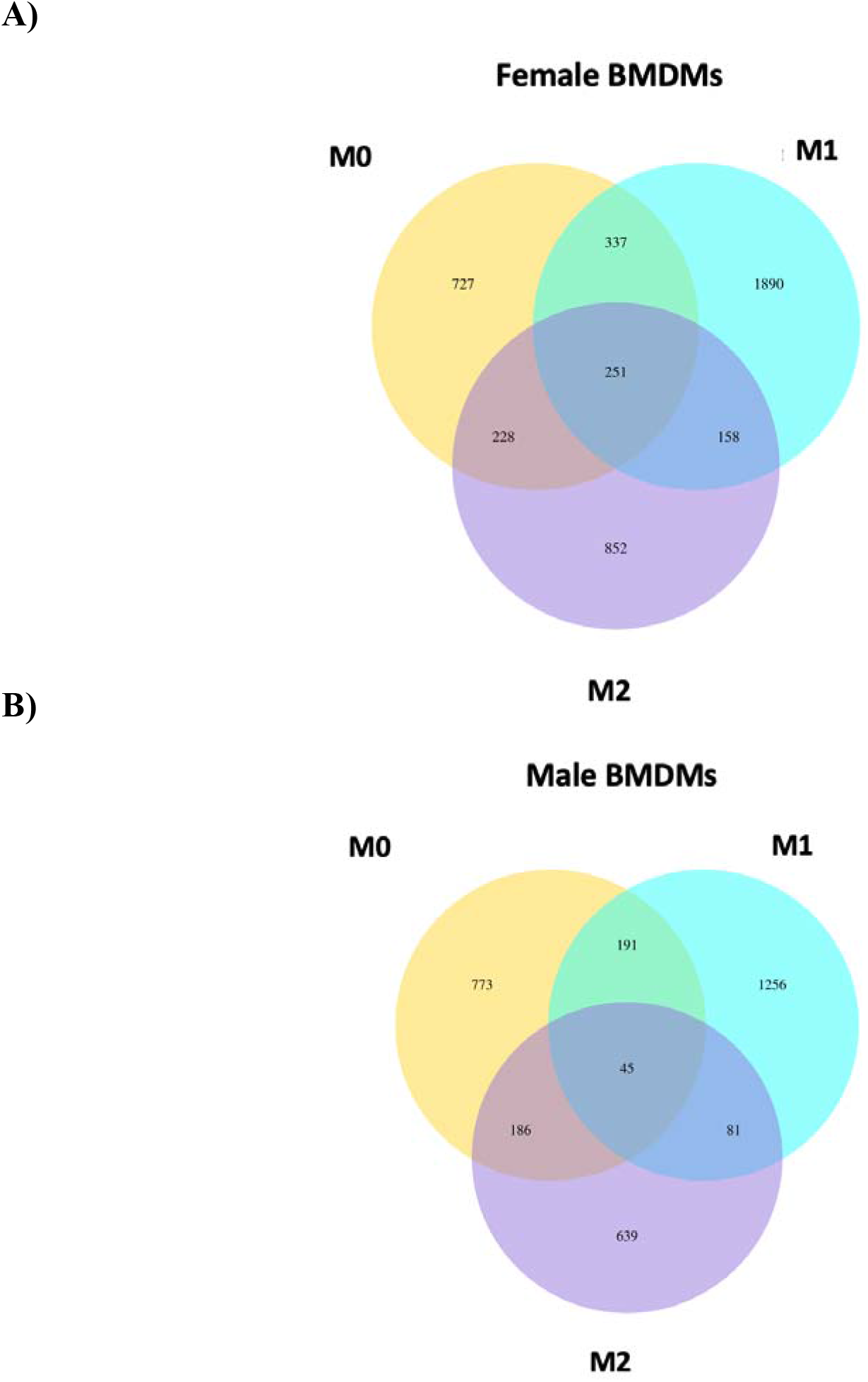
Venn diagrams showing numbers of shared and unique DEGs induced by arsenic exposure across stimulation groups in female and male BMDMs. Venn Diagrams were generated with our DEG cut-off |log2FC|>0 and adjusted *p*-value of <0.05 for Female (A) and Male (B) groups. For both diagrams, yellow circles contain DEGs in the M0 stimulation group, aqua circles in the M1 stimulation group, and purple circles in the M2 stimulation group.

### STRING Protein-Protein Networks of iAs-exposed BMDMs

STRING (Shannon 2003) automatically provides information regarding each protein-protein interaction (PPI) network, including the number of nodes (before disconnecting), the number of edges vs. expected number of edges, the average node degree, the average clustering coefficient, and the PPI enrichment p-value (**Table 2-14**). Given our input criteria, networks were generated for all female-derived cells except for the M2 downregulated gene set (**Fig. 2-4**). Considering the PPI enrichment *p-*values, all female networks that were generated except for the M0 downregulated set had statistically significant interaction scores (*p*<0.05) (**Table 2-14**). For the male-derived cells, STRING networks were generated for the M0 upregulated, M1 down- and upregulated, and M2 upregulated groups (**Fig. 2-4**); however, considering the PPI enrichment *p-*value only the male M2 unregulated gene set had a statistically significant interaction score (*p*<0.05) (**Table 2-14**).

**Figure 2-4.**
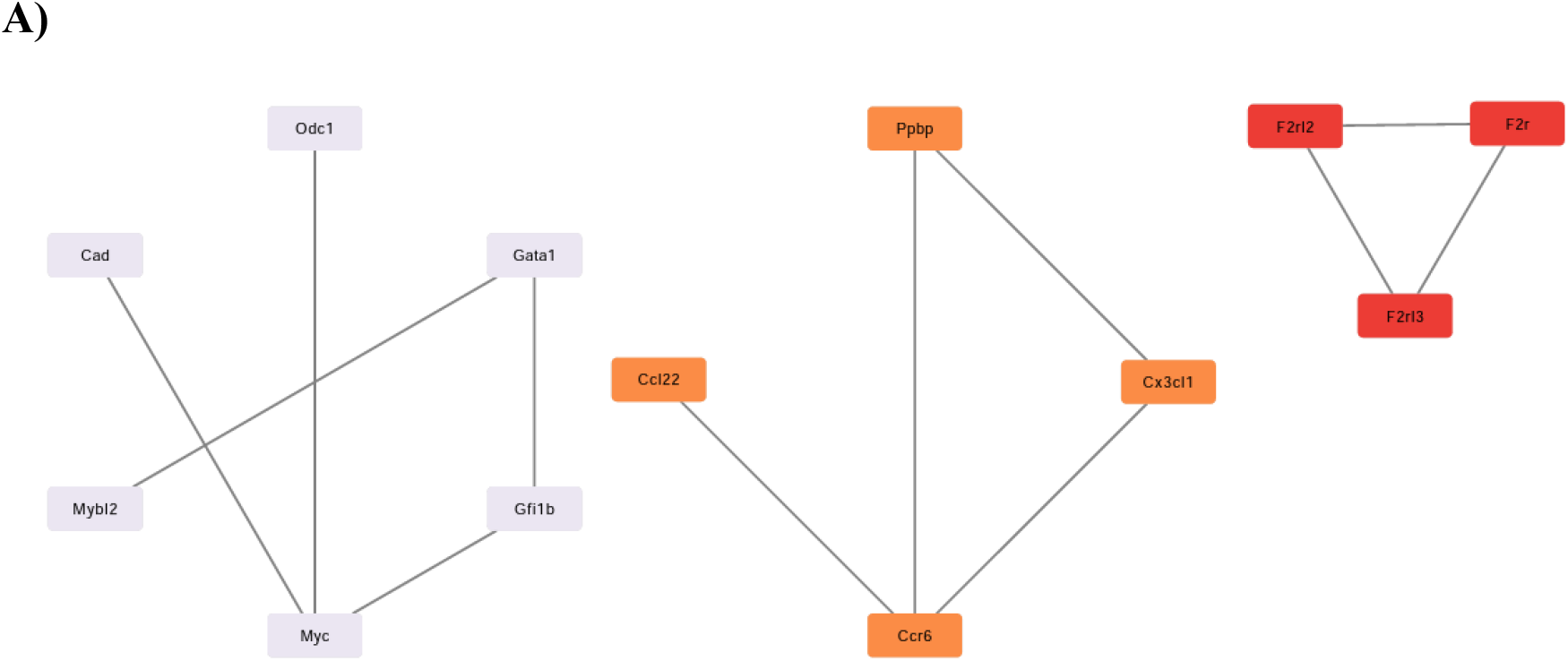
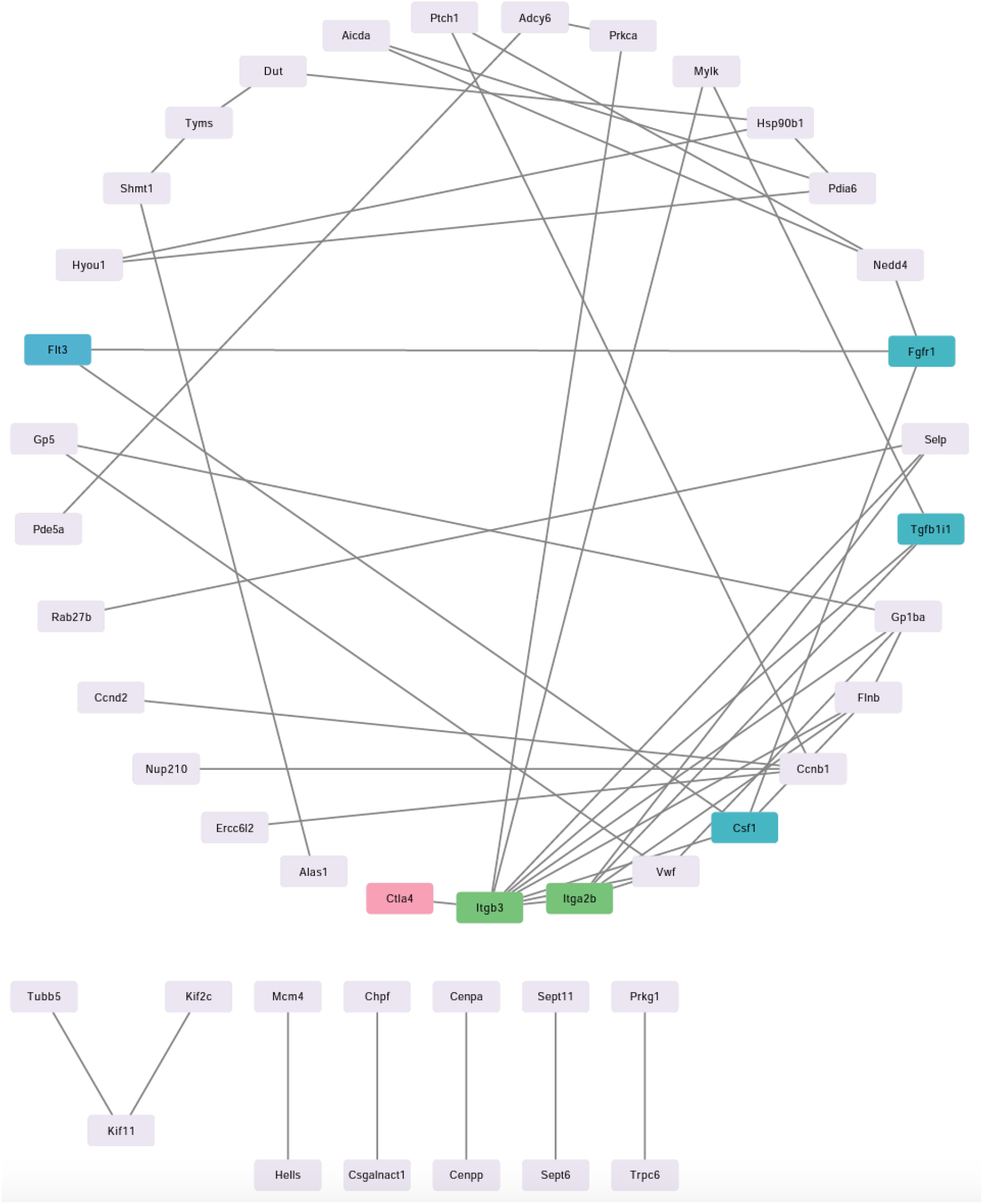
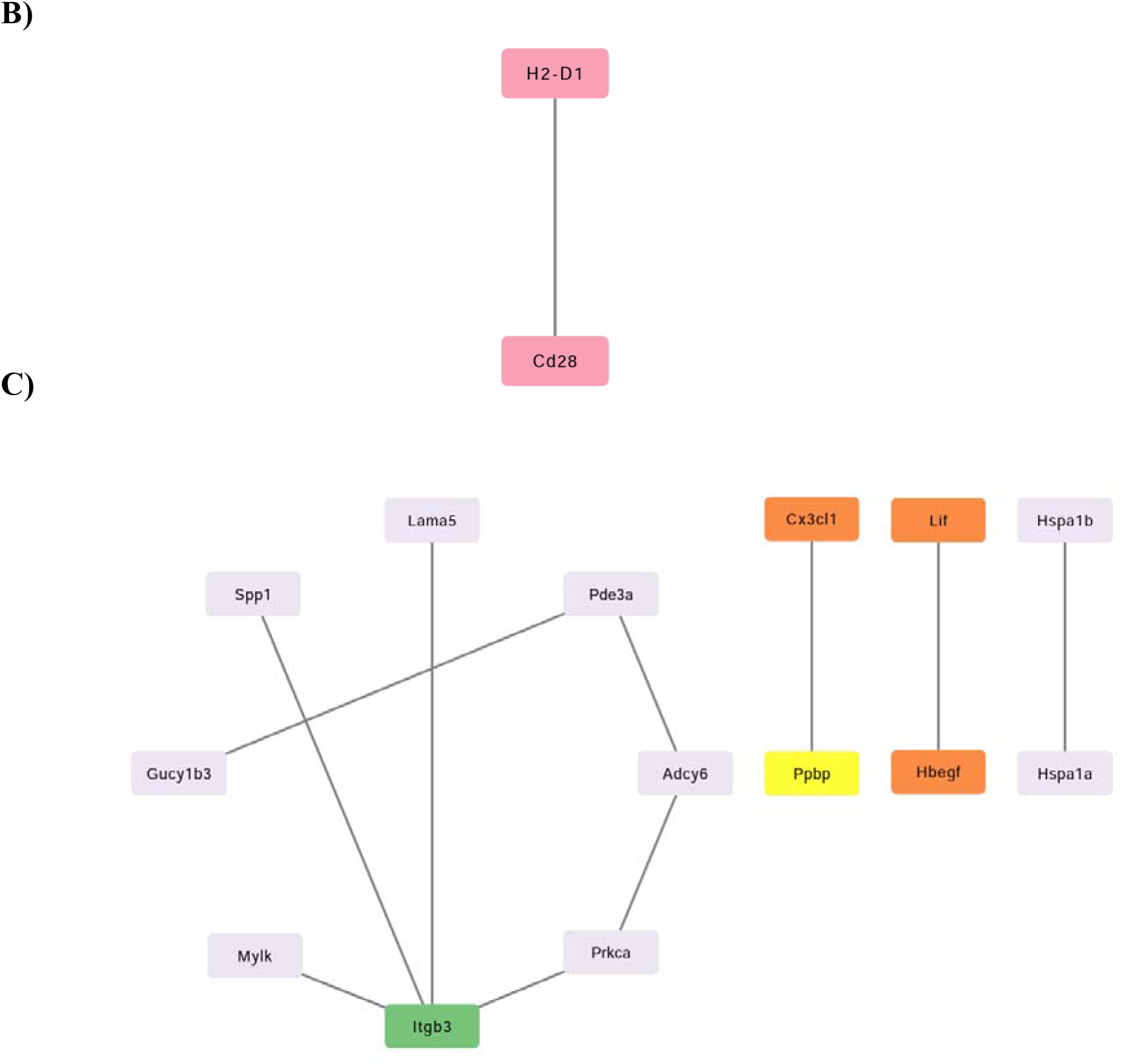
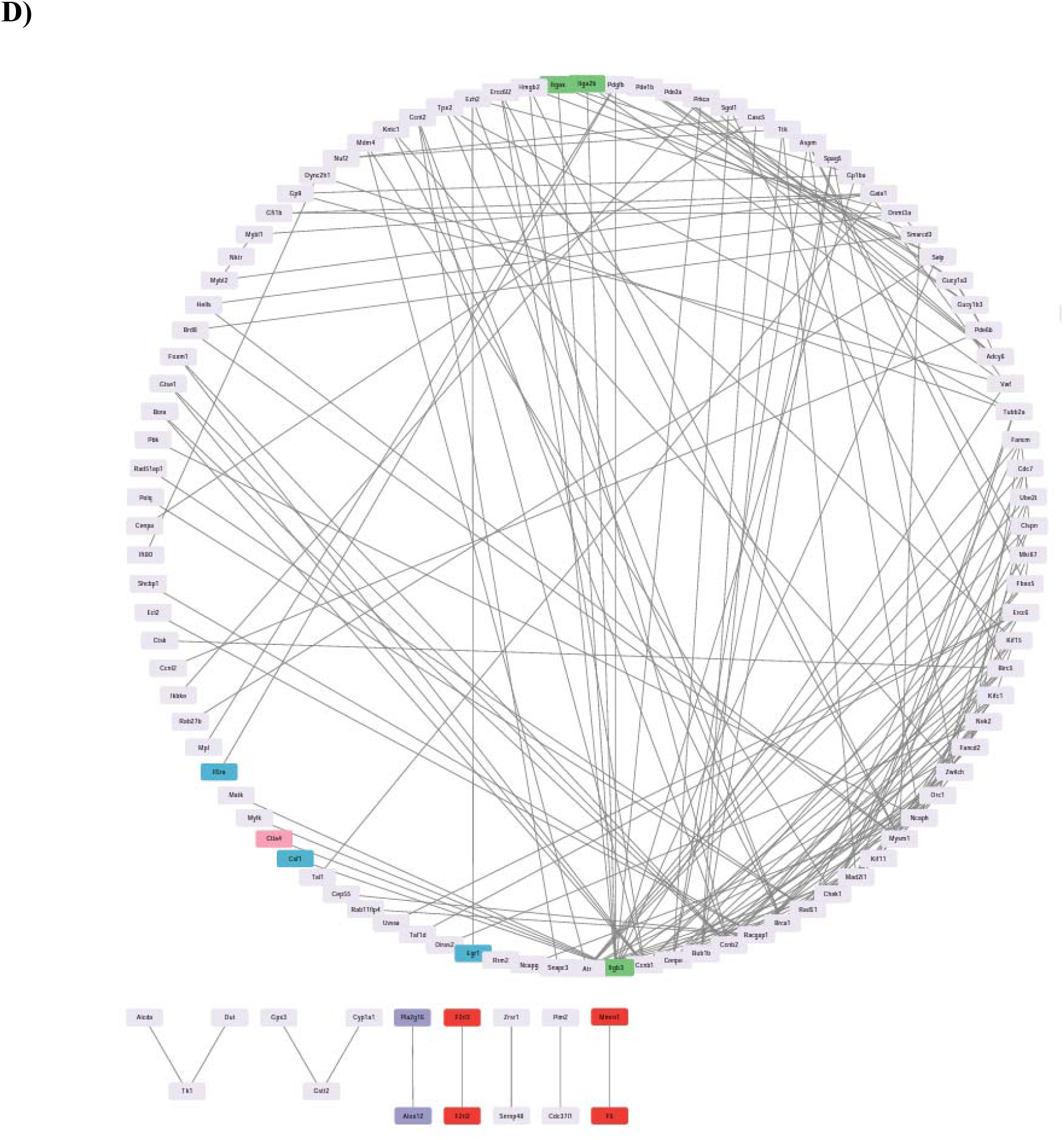
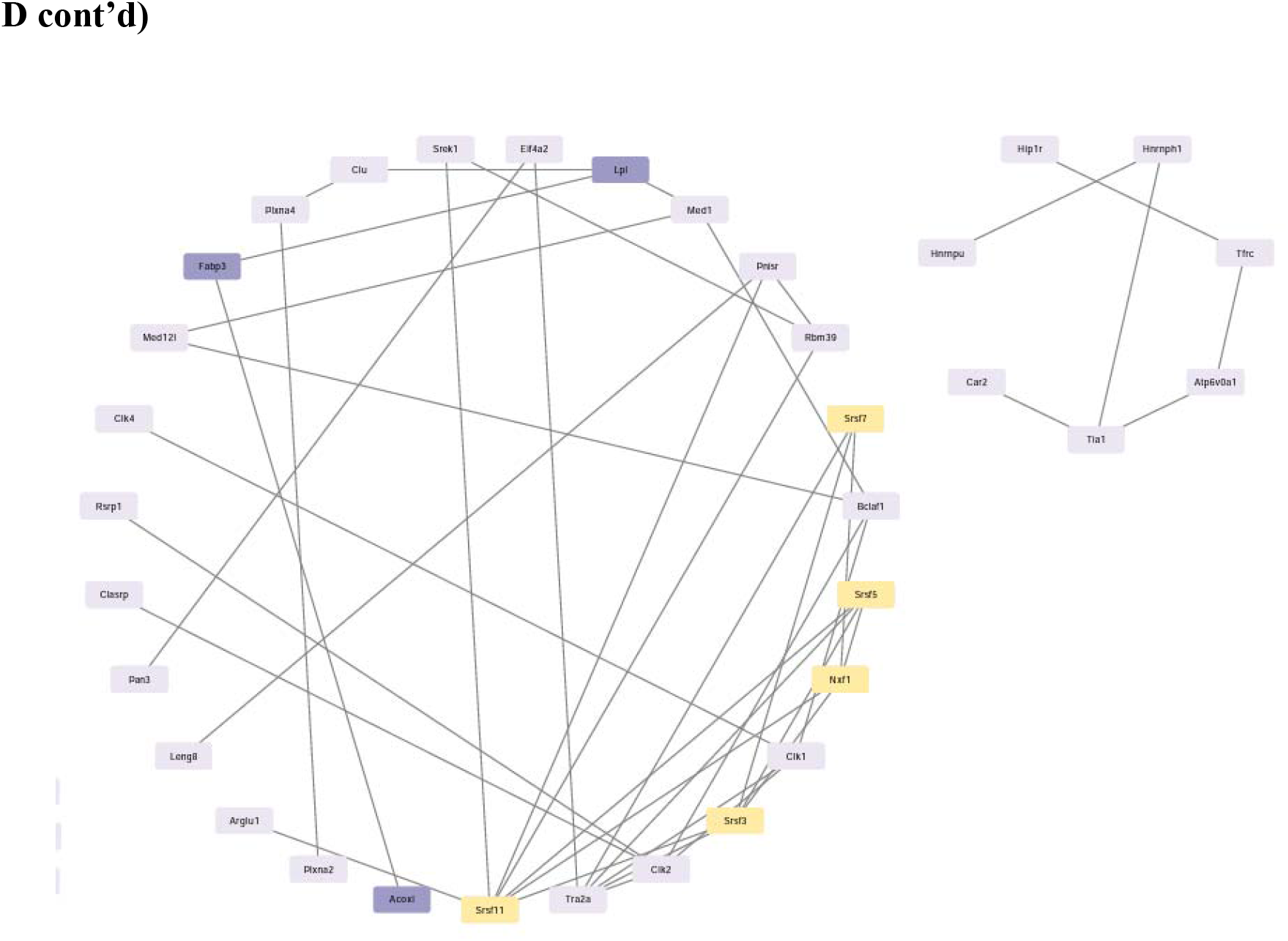
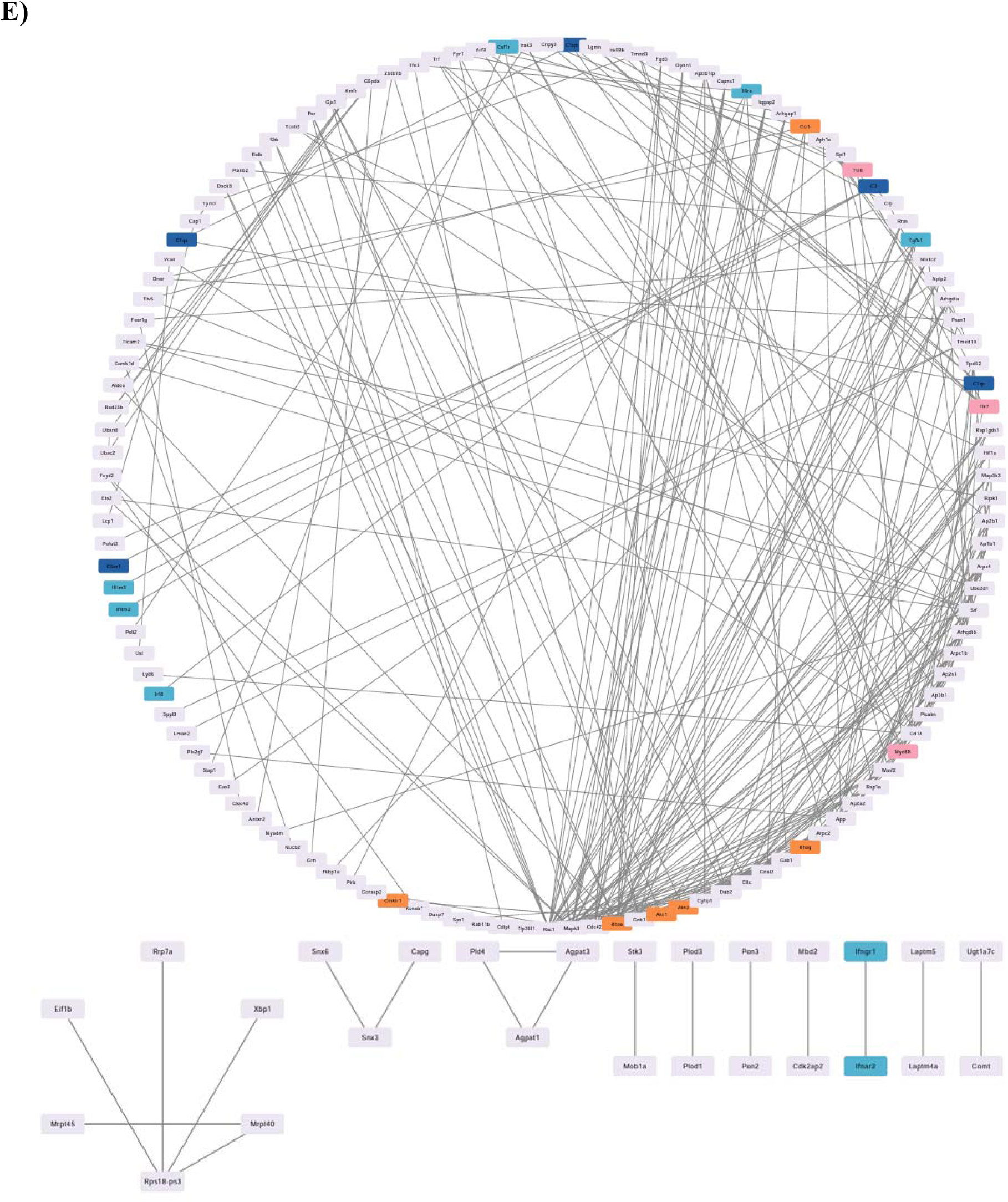
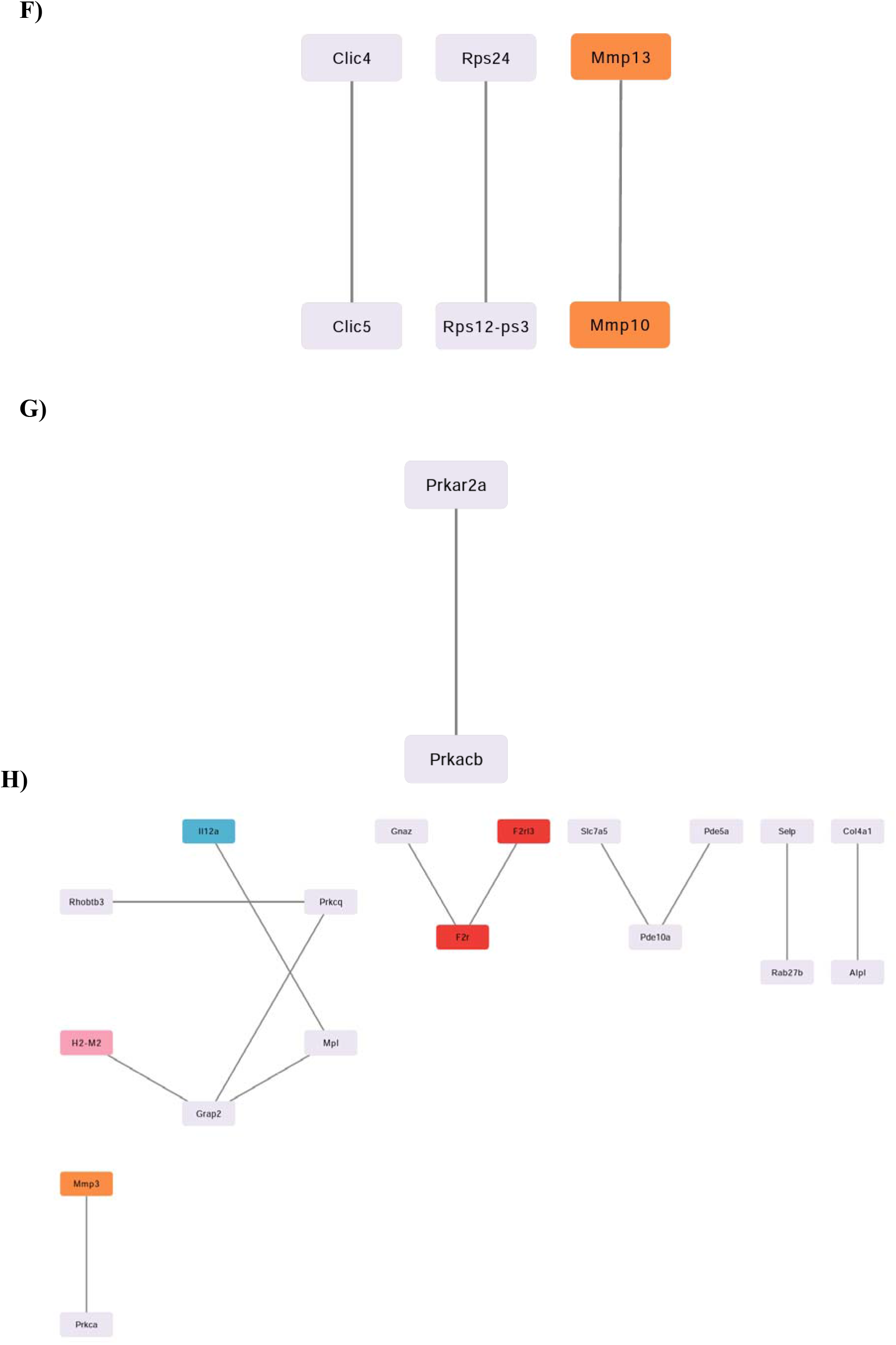
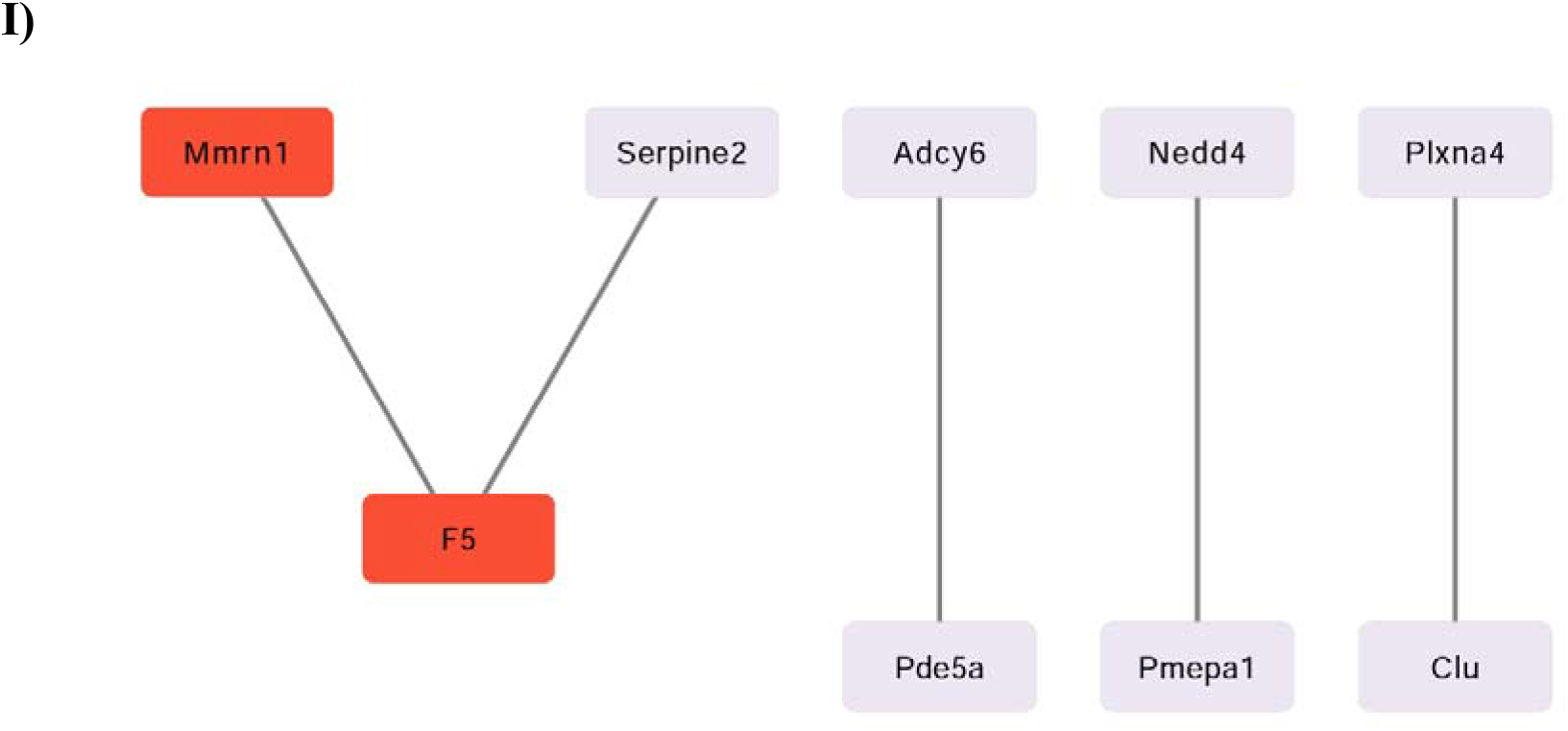
Cytoscape maps of STRING PPI networks for up- and downregulated genes from female and male iAs-exposed BMDMs across stimulation groups. Maps of PPI networks were generated in STRING and visualized in Cytoscape for Female M0 up-(A) and downregulated (B), Male M0 upregulated (C), Female M1 up-(D) and downregulated (E), Male M1 up-(F) and downregulated (G), Female M2 upregulated (H), and Male M2 upregulated (I) groups. Nodes are color coded according to immune relevant categories: integrin signaling (green), coagulation factors (red), chemotaxis and chemokine signaling (orange), pattern recognition and antigen presentation (pink), splicing machinery (yellow), lipid metabolism (purple), complement system (blue), and cytokine signaling (aqua). If no map was generated for a comparison group, that group is omitted from this figure.

**Table 2-2.** Top 50 upregulated genes in iAs-treated M0 female-derived BMDMs. If fewer than 50 genes met the more stringent cutoff of |log2FC| > 0.2, only those genes are listed.

| Gene name | Description | log2FC | p-adj | Chromosome |
| --- | --- | --- | --- | --- |
| Dlk1 | delta like non-canonical Notch ligand 1 | 6.33 | 1.88E-06 | 12 |
| Gm13994 | predicted gene 13994 | 5.08 | 1.55E-07 | 2 |
| Igkv1-99 | immunoglobulin kappa variable 1-99 | 4.62 | 0.00018138 | 6 |
| Ccl22 | chemokine (C-C motif) ligand 22 | 3.52 | 2.46E-12 | 8 |
| Ctla4 | cytotoxic T-lymphocyte-associated protein 4 | 3.26 | 1.59E-07 | 1 |
| Timd2 | T cell immunoglobulin and mucin domain containing 2 | 2.83 | 0.01844349 | 11 |
| Zbtb32 | zinc finger and BTB domain containing 32 | 2.64 | 7.67E-10 | 7 |
| Cyp1a1 | cytochrome P450, family 1, subfamily a, polypeptide 1 | 2.61 | 6.78E-16 | 9 |
| Ighv7-3 | immunoglobulin heavy variable 7-3 | 2.41 | 2.67E-12 | 12 |
| Cdh17 | cadherin 17 | 2.38 | 9.64E-06 | 4 |
| Shisa8 | shisa family member 8 | 2.19 | 1.07E-10 | 15 |
| Aicda | activation-induced cytidine deaminase | 2.17 | 0.00013317 | 6 |
| Ripor3 | RIPOR family member 3 | 1.94 | 0.04701391 | 2 |
| Alpl | alkaline phosphatase, liver/bone/kidney | 1.89 | 0.04701391 | 4 |
| Grm6 | glutamate receptor, metabotropic 6 | 1.87 | 0.01532506 | 11 |
| Mef2b | myocyte enhancer factor 2B | 1.84 | 1.55E-07 | 8 |
| Igkv14-126 | immunoglobulin kappa variable 14-126 | 1.76 | 0.00619194 | 6 |
| Pdcd1 | programmed cell death 1 | 1.76 | 1.84E-05 | 1 |
| Ighv10-1 | immunoglobulin heavy variable 10-1 | 1.7 | 4.33E-05 | 12 |
| Igkv8-30 | immunoglobulin kappa chain variable 8-30 | 1.68 | 0.00144098 | 6 |
| Mcts2 | malignant T cell amplified sequence 2 | 1.58 | 0.01978746 | 2 |
| Il5ra | interleukin 5 receptor, alpha | 1.51 | 5.73E-07 | 6 |
| Cenpp | centromere protein P | 1.51 | 0.03031702 | 13 |
| St8sia6 | ST8 alpha-N-acetyl-neuraminide alpha-2,8-sialyltransferase 6 | 1.48 | 1.16E-07 | 2 |
| Serpina3b | serine (or cysteine) peptidase inhibitor, clade A, member 3B | 1.46 | 0.00135878 | 12 |
| Ankrd35 | ankyrin repeat domain 35 | 1.45 | 0.02928146 | 3 |
| Sec14l2 | SEC14-like lipid binding 2 | 1.42 | 0.00055465 | 11 |
| Flt3 | FMS-like tyrosine kinase 3 | 1.37 | 0.01327819 | 5 |
| Prkg1 | protein kinase, cGMP-dependent, type I | 1.37 | 0.0301031 | 19 |
| Serpinb12 | serine (or cysteine) peptidase inhibitor, clade B (ovalbumin), member 12 | 1.33 | 0.02199822 | 1 |
| Hic1 | hypermethylated in cancer 1 | 1.31 | 2.05E-05 | 11 |
| Selp | selectin, platelet | 1.29 | 0.00094835 | 1 |
| Amigo2 | adhesion molecule with Ig like domain 2 | 1.28 | 0.00063432 | 15 |
| Cx3cl1 | chemokine (C-X3-C motif) ligand 1 | 1.26 | 0.01368144 | 8 |
| Dapk2 | death-associated protein kinase 2 | 1.22 | 1.84E-05 | 9 |
| Tesc | tescalcin | 1.22 | 0.00093705 | 5 |
| Ccdc8 | coiled-coil domain containing 8 | 1.21 | 0.00135878 | 7 |
| Lrig1 | leucine-rich repeats and immunoglobulin-like domains 1 | 1.18 | 0.02178561 | 6 |
| Tgfbli1 | transforming growth factor beta 1 induced transcript 1 | 1.18 | 0.01153451 | 7 |
| F2rl3 | coagulation factor II (thrombin) receptor-like 3 | 1.17 | 0.04165778 | 8 |
| Ache | acetylcholinesterase | 1.17 | 0.01834041 | 5 |
| Ccr6 | chemokine (C-C motif) receptor 6 | 1.16 | 0.00017717 | 17 |
| Fcrl5 | Fc receptor-like 5 | 1.15 | 0.00019303 | 3 |
| Nek2 | NIMA (never in mitosis gene a)-related expressed kinase 2 | 1.14 | 0.00083141 | 1 |
| Gp5 | glycoprotein 5 (platelet) | 1.13 | 1.12E-05 | 16 |
| Fah | fumarylacetoacetate hydrolase | 1.13 | 0.04640585 | 7 |
| F7 | coagulation factor VII | 1.12 | 5.73E-07 | 8 |
| Mmrn1 | multimerin 1 | 1.08 | 0.00237456 | 6 |
| Cavin2 | caveolae associated 2 | 1.08 | 0.0004847 | 1 |
| Lrrc32 | leucine rich repeat containing 32 | 1.05 | 0.00147816 | 7 |

**Table 2-3.** Top 50 downregulated genes in iAs-treated M0 female-derived BMDMs. If fewer than 50 genes met the more stringent cutoff of |log2FC| > 0.2, only those genes are listed.

| Gene name | Description | log2FC | p-adj | Chromosome |
| --- | --- | --- | --- | --- |
| Fcrls | Fc receptor-like S, scavenger receptor | -1.49 | 4.66E-11 | 3 |
| Dynlt1a | dynein light chain Tctex-type 1A | -1.14 | 0.00172736 | 17 |
| Fxyd2 | FXDY domain-containing ion transport regulator 2 | -1.07 | 9.53E-07 | 9 |
| Gm5960 | predicted gene 5960 | -0.95 | 0.00863037 | 15 |
| Sirpb1a | signal-regulatory protein beta 1A | -0.81 | 0.00412515 | 3 |
| I830127L07Rik | RIKEN cDNA I830127L07 gene | -0.67 | 0.03950975 | 15 |
| Ifi2712a | interferon, alpha-inducible protein 27 like 2A | -0.58 | 0.03468125 | 12 |
| Cd28 | CD28 antigen | -0.56 | 0.04646198 | 1 |
| Ncapg2 | non-SMC condensin II complex, subunit G2 | -0.49 | 3.64E-06 | 12 |
| 5430427O19Rik | RIKEN cDNA 5430427O19 gene | -0.42 | 0.01217715 | X |
| Clec4a3 | C-type lectin domain family 4, member a3 | -0.41 | 0.03560751 | 6 |
| Siglec1 | sialic acid binding Ig-like lectin 1, sialoadhesin | -0.41 | 0.03136071 | 2 |
| TLR7 | toll-like receptor 7 | -0.39 | 0.00247724 | X |
| Bcl6 | B cell leukemia/lymphoma 6 | -0.39 | 0.00319075 | 16 |
| Crybg1 | crystallin beta-gamma domain containing 1 | -0.35 | 0.04753275 | 10 |
| Tbkbp1 | TBK1 binding protein 1 | -0.33 | 0.03448038 | 11 |
| Ifi207 | interferon activated gene 207 | -0.32 | 0.0322534 | 1 |
| S1pr1 | sphingosine-1-phosphate receptor 1 | -0.32 | 0.04343788 | 3 |
| H2-D1 | histocompatibility 2, D region locus 1 | -0.31 | 0.02475833 | 17 |

| Gene name | Description | log2FC | *p*-adj | Chromosome |
| --- | --- | --- | --- | --- |
| Rassf2 | Ras association<br>(RalGDS/AF-6) domain<br>family member 2 | -0.31 | 0.04808839 | 2 |
| Lpin2 | lipin 2 | -0.31 | 0.00629847 | 17 |
| Stom | stomatin | -0.3 | 0.03468125 | 2 |
| Lpxn | leupaxin | -0.3 | 0.04494512 | 19 |
| Sec14l1 | SEC14-like lipid binding 1 | -0.28 | 0.01844349 | 11 |
| Zbtb7b | zinc finger and BTB<br>domain containing 7B | -0.27 | 0.0278719 | 3 |
| Irf8 | interferon regulatory factor<br>8 | -0.25 | 0.0297214 | 8 |
| Itm2b | integral membrane protein<br>2B | -0.24 | 0.01154089 | 14 |

**Table 2-4.**
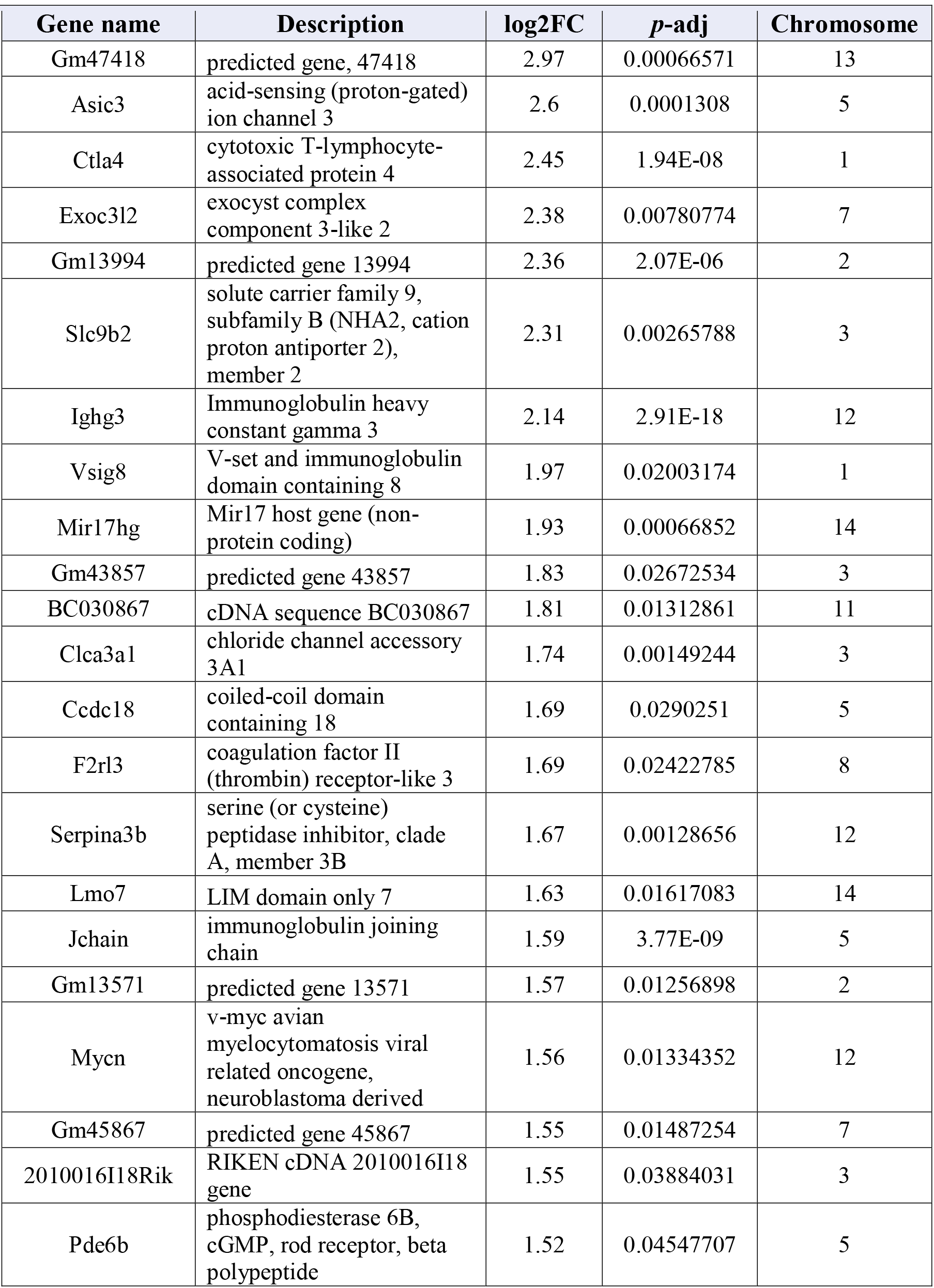
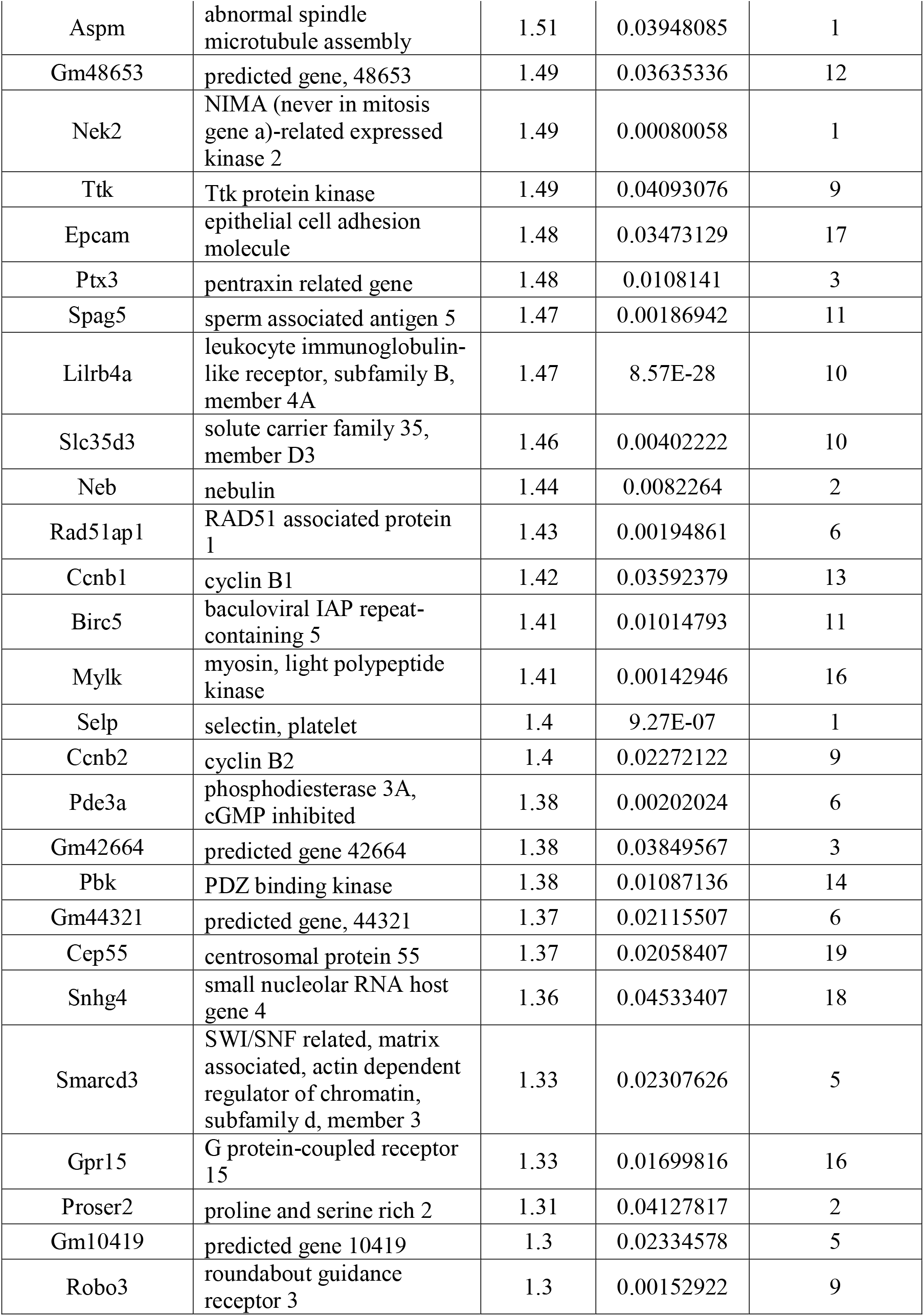
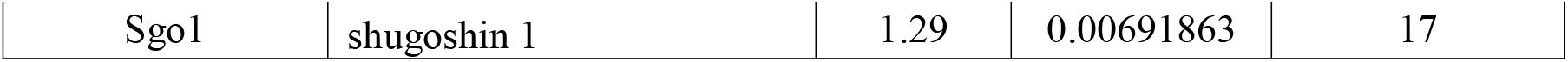
Top 50 upregulated genes in iAs-treated M1 female-derived BMDMs. If fewer than 50 genes met the more stringent cutoff of |log2FC| > 0.2, only those genes are listed.

**Table 2-5.** Top 50 downregulated genes in iAs-treated M1 female-derived BMDMs. If fewer than 50 genes met the more stringent cutoff of |log2FC| > 0.2, only those genes are listed.

| Gene name | Description | log2FC | p-adj | Chromosome |
| --- | --- | --- | --- | --- |
| BC100530 | cDNA sequence BC100530 | -1.69 | 0.04146252 | 16 |
| Stfa1 | stefin A1 | -1.63 | 0.00619013 | 16 |
| Dner | delta/notch-like EGF repeat containing | -1.49 | 0.01980947 | 1 |
| Fxyd2 | FXYD domain-containing ion transport regulator 2 | -1.23 | 0.00036899 | 9 |
| Rps18-ps3 | ribosomal protein S18, pseudogene 3 | -1.21 | 0.03808157 | 8 |
| 2010005H15Rik | RIKEN cDNA 2010005H15 gene | -1.08 | 0.03571232 | 16 |
| Ifitm6 | interferon induced transmembrane protein 6 | -1.05 | 0.00056282 | 7 |
| Trf | transferrin | -0.97 | 0.01796125 | 9 |
| Lancl3 | LanC lantibiotic synthetase component C-like 3 (bacterial) | -0.94 | 0.04950367 | X |
| Stfa3 | stefin A3 | -0.93 | 0.02420328 | 16 |
| Spint1 | serine protease inhibitor, Kunitz type 1 | -0.86 | 0.03662405 | 2 |
| Fcrls | Fc receptor-like S, scavenger receptor | -0.86 | 0.00028693 | 3 |
| Gja1 | gap junction protein, alpha 1 | -0.85 | 1.04E-05 | 10 |
| Mmp8 | matrix metalloproteinase 8 | -0.82 | 0.00038347 | 9 |
| Gm13736 | predicted gene 13736 | -0.77 | 0.00618654 | 2 |
| Rcbtb2 | regulator of chromosome condensation (RCC1) and BTB (POZ) domain containing protein 2 | -0.75 | 0.01886835 | 14 |
| Cd36 | CD36 molecule | -0.73 | 0.00015923 | 5 |
| Gm17586 | predicted gene, 17586 | -0.73 | 0.00545644 | 11 |
| Vcan | versican | -0.71 | 0.02534998 | 13 |
| Chil3 | chitinase-like 3 | -0.67 | 0.02716435 | 3 |
| Ifitm2 | interferon induced transmembrane protein 2 | -0.66 | 5.37E-05 | 7 |
| Sirpb1a | signal-regulatory protein beta 1A | -0.66 | 0.00614589 | 3 |
| Zdhhc14 | zinc finger, DHHC domain containing 14 | -0.65 | 0.01773723 | 17 |
| Gm5150 | predicted gene 5150 | -0.64 | 0.01710856 | 3 |
| Clec4a2 | C-type lectin domain family 4, member a2 | -0.64 | 0.00423153 | 6 |
| Shb | src homology 2 domain-containing transforming protein B | -0.64 | 0.00664065 | 4 |
| Tifab | TRAF-interacting protein with forkhead-associated domain, family member B | -0.64 | 0.00736189 | 13 |
| Peli2 | pellino 2 | -0.6 | 0.01710856 | 14 |
| Klra2 | killer cell lectin-like receptor, subfamily A, member 2 | -0.59 | 0.00108968 | 6 |
| Clec4a1 | C-type lectin domain family 4, member a1 | -0.59 | 1.92E-09 | 6 |
| Mmp27 | matrix metalloproteinase 27 | -0.59 | 0.0436478 | 9 |
| C1qa | complement component 1, q subcomponent, alpha polypeptide | -0.58 | 0.00572425 | 4 |
| Dab2 | disabled 2, mitogen-responsive phosphoprotein | -0.57 | 0.00216923 | 15 |
| Gpr141 | G protein-coupled receptor 141 | -0.57 | 0.00207595 | 13 |
| Ccr5 | chemokine (C-C motif) receptor 5 | -0.57 | 0.01886835 | 9 |
| Snta1 | syntrophin, acidic 1 | -0.56 | 0.00182223 | 2 |
| Emilin2 | elastin microfibril interfacer 2 | -0.55 | 0.0201013 | 17 |
| Sirpb1b | signal-regulatory protein beta 1B | -0.55 | 0.03925105 | 3 |
| Siglec1 | sialic acid binding Ig-like lectin 1, sialoadhesin | -0.55 | 0.00052449 | 2 |
| C130050O18Rik | RIKEN cDNA C130050O18 gene | -0.54 | 0.01811698 | 5 |
| Syn1 | synapsin I | -0.54 | 0.013775 | X |
| Ust | uronyl-2-sulfotransferase | -0.54 | 0.00977457 | 10 |
| Pon3 | paraoxonase 3 | -0.53 | 0.00037048 | 6 |
| Sirpb1c | signal-regulatory protein beta 1C | -0.53 | 0.01761956 | 3 |
| Ophn1 | oligophrenin 1 | -0.53 | 0.00889486 | X |
| Adgrl2 | adhesion G protein-coupled receptor L2 | -0.52 | 0.03731865 | 3 |
| Tlr8 | toll-like receptor 8 | -0.52 | 0.00937413 | X |
| C5ar1 | complement component 5a receptor 1 | -0.52 | 0.00107572 | 7 |
| Serpinb8 | serine (or cysteine) peptidase inhibitor, clade B, member 8 | -0.52 | 0.00222194 | 1 |
| Nfatc2 | nuclear factor of activated T cells, cytoplasmic, calcineurin dependent 2 | -0.52 | 0.00186942 | 2 |

**Table 2-6.** Top 50 upregulated genes in iAs-treated M2 female-derived BMDMs. If fewer than 50 genes met the more stringent cutoff of |log2FC| > 0.2, only those genes are listed.

| Gene name | Description | log2FC | p-adj | Chromosome |
| --- | --- | --- | --- | --- |
| Ighv3-8 | immunoglobulin heavy variable V3-8 | 3.01 | 0.02513771 | 12 |
| Sox5 | SRY (sex determining region Y)-box 5 | 2.61 | 0.02962507 | 6 |
| Alpl | alkaline phosphatase, liver/bone/kidney | 2.37 | 0.00010936 | 4 |
| Lmo7 | LIM domain only 7 | 2.32 | 0.00573742 | 14 |
| Prnt4 | proline-rich transmembrane protein 4 | 2.12 | 0.00177232 | 6 |
| Igkv8-21 | immunoglobulin kappa variable 8-21 | 2.12 | 0.02554368 | 6 |
| Fah | fumarylacetoacetate hydrolase | 2.11 | 0.03806948 | 7 |
| Timd2 | T cell immunoglobulin and mucin domain containing 2 | 2.08 | 0.02104574 | 11 |
| Cyp1a1 | cytochrome P450, family 1, subfamily a, polypeptide 1 | 2.03 | 3.54E-08 | 9 |
| Dkk3 | dickkopf WNT signaling pathway inhibitor 3 | 2.01 | 0.01640196 | 7 |
| Ripor3 | RIPOR family member 3 | 1.92 | 0.01528344 | 2 |
| Trim58 | tripartite motif-containing 58 | 1.8 | 0.02888198 | 11 |
| St8sia6 | ST8 alpha-N-acetylneuraminide alpha-2,8-sialyltransferase 6 | 1.66 | 0.003113 | 2 |
| Kcnk5 | potassium channel, subfamily K, member 5 | 1.65 | 0.00088943 | 14 |
| Pde10a | phosphodiesterase 10A | 1.58 | 0.0060877 | 17 |
| Il12a | interleukin 12a | 1.55 | 0.0228665 | 3 |
| Sla2 | Src-like-adaptor 2 | 1.54 | 0.00441153 | 2 |
| Igkv1-110 | immunoglobulin kappa variable 1-110 | 1.53 | 0.01615671 | 6 |
| Serpina3b | serine (or cysteine) peptidase inhibitor, clade A, member 3B | 1.48 | 0.00441153 | 12 |
| Amigo2 | adhesion molecule with Ig like domain 2 | 1.46 | 0.03263407 | 15 |
| Srgap3 | SLIT-ROBO Rho GTPase activating protein 3 | 1.37 | 0.02888198 | 6 |
| Selp | selectin, platelet | 1.36 | 0.00025267 | 1 |
| Mrvi1 | MRV integration site 1 | 1.34 | 0.03806948 | 7 |
| Slc14a1 | solute carrier family 14 (urea transporter), member 1 | 1.32 | 1.12E-12 | 18 |
| Mpl | myeloproliferative leukemia virus oncogene | 1.31 | 0.02104574 | 4 |
| Rpl17 | ribosomal protein L17 | 1.3 | 7.63E-06 | 18 |
| Cavin2 | caveolae associated 2 | 1.29 | 0.01022192 | 1 |
| F2rl3 | coagulation factor II (thrombin) receptor-like 3 | 1.28 | 0.04635178 | 8 |
| Gfi1b | growth factor independent 1B | 1.23 | 0.0028452 | 2 |
| Kcnj5 | potassium inwardly-rectifying channel, subfamily J, member 5 | 1.22 | 0.04111068 | 9 |
| Samd14 | sterile alpha motif domain containing 14 | 1.2 | 0.03692125 | 11 |
| Mylk | myosin, light polypeptide kinase | 1.17 | 0.00151765 | 16 |
| Dusp10 | dual specificity phosphatase 10 | 1.16 | 0.01395409 | 1 |
| Rhobtb3 | Rho-related BTB domain containing 3 | 1.12 | 0.03806948 | 13 |
| Rab27b | RAB27B, member RAS oncogene family | 1.1 | 0.00177232 | 18 |
| Gnaz | guanine nucleotide binding protein, alpha z subunit | 1.08 | 0.03936691 | 10 |
| Pcyt1b | phosphate cytidyltransferase 1, choline, beta isoform | 1.06 | 0.03263407 | X |
| Rbpms2 | RNA binding protein with multiple splicing 2 | 1.06 | 0.03951399 | 9 |
| Prkca | protein kinase C, alpha | 1.06 | 8.91E-05 | 11 |
| Trpc6 | transient receptor potential cation channel, subfamily C, member 6 | 1.02 | 0.03806948 | 9 |
| F2r | coagulation factor II (thrombin) receptor | 1.01 | 5.12E-06 | 13 |
| F11r | F11 receptor | 0.93 | 0.03029747 | 1 |
| Ppic | peptidylprolyl isomerase C | 0.93 | 0.01688191 | 18 |
| Pde5a | phosphodiesterase 5A,<br>cGMP-specific | 0.92 | 0.003113 | 3 |
| Pdlim1 | PDZ and LIM domain 1<br>(elfin) | 0.92 | 0.02813309 | 19 |
| Prkcq | protein kinase C, theta | 0.92 | 0.00278799 | 2 |
| Zfp503 | zinc finger protein 503 | 0.91 | 0.00163697 | 14 |
| Fads2 | fatty acid desaturase 2 | 0.9 | 0.00694781 | 19 |
| Col4a1 | collagen, type IV, alpha 1 | 0.9 | 0.01606158 | 8 |
| Rab11fip4 | RAB11 family interacting<br>protein 4 (class II) | 0.89 | 0.00041913 | 11 |

**Table 2-7.** Top 50 downregulated genes in iAs-treated M2 female-derived BMDMs. If fewer than 50 genes met the more stringent cutoff of |log2FC| > 0.2, only those genes are listed.

| Gene name | Description | log2FC | p-adj | Chromosome |
| --- | --- | --- | --- | --- |
| Retnla | resistin like alpha | -1.42 | 0.04418227 | 16 |
| Agmo | alkylglycerol monooxygenase | -1.14 | 0.0025286 | 12 |
| Lrg1 | leucine-rich alpha-2-glycoprotein 1 | -1.11 | 0.00304106 | 17 |
| Prom1 | prominin 1 | -1.08 | 0.01688191 | 5 |
| Stfa2l1 | stefin A2 like 1 | -0.95 | 0.01188261 | 16 |
| Trf | transferrin | -0.93 | 0.00278799 | 9 |
| Slc9a9 | solute carrier family 9 (sodium/hydrogen exchanger), member 9 | -0.87 | 7.58E-12 | 9 |
| Slc46a3 | solute carrier family 46, member 3 | -0.86 | 0.01395409 | 5 |
| Cldn1 | claudin 1 | -0.85 | 0.02101633 | 16 |
| Rnase4 | ribonuclease, RNase A family 4 | -0.85 | 0.00201385 | 14 |
| Abca9 | ATP-binding cassette, sub-family A (ABC1), member 9 | -0.82 | 6.01E-05 | 11 |
| Hsd11b1 | hydroxysteroid 11-beta dehydrogenase 1 | -0.76 | 0.02803637 | 1 |
| Chil3 | chitinase-like 3 | -0.74 | 0.00068731 | 3 |
| Ccr2 | chemokine (C-C motif) receptor 2 | -0.74 | 0.01369382 | 9 |
| Tbc1d4 | TBC1 domain family, member 4 | -0.69 | 0.01664389 | 14 |
| Slco2b1 | solute carrier organic anion transporter family, member 2b1 | -0.67 | 0.01664389 | 7 |
| Cd28 | CD28 antigen | -0.65 | 0.03164287 | 1 |
| Smagp | small cell adhesion glycoprotein | -0.63 | 0.00758715 | 15 |
| Clec4a1 | C-type lectin domain family 4, member a1 | -0.61 | 0.01441115 | 6 |
| Wwp1 | WW domain containing E3 ubiquitin protein ligase 1 | -0.58 | 7.63E-06 | 4 |
| Rcbtb2 | regulator of chromosome condensation (RCC1) and BTB (POZ) domain containing protein 2 | -0.58 | 0.01295793 | 14 |
| Sh3d19 | SH3 domain protein D19 | -0.56 | 0.00607446 | 3 |
| Slc16a7 | solute carrier family 16 (monocarboxylic acid transporters), member 7 | -0.55 | 0.03191033 | 10 |
| Ly6c2 | lymphocyte antigen 6 complex, locus C2 | -0.54 | 0.03163196 | 15 |
| Clec10a | C-type lectin domain family 10, member A | -0.51 | 0.03819733 | 11 |
| P2ry6 | pyrimidinergic receptor P2Y, G-protein coupled, 6 | -0.45 | 0.04635178 | 7 |
| Hpse | heparanase | -0.42 | 0.00391528 | 5 |
| Rnf150 | ring finger protein 150 | -0.41 | 0.00033217 | 8 |
| Pygl | liver glycogen phosphorylase | -0.4 | 2.02E-05 | 12 |
| Vsir | V-set immunoregulatory receptor | -0.39 | 0.00088943 | 10 |
| Gda | guanine deaminase | -0.39 | 0.00278799 | 19 |
| P2rx7 | purinergic receptor P2X, ligand-gated ion channel, 7 | -0.38 | 0.02888198 | 5 |
| Rab11fip5 | RAB11 family interacting protein 5 (class I) | -0.37 | 0.03806948 | 6 |
| Nfatc2 | nuclear factor of activated T cells, cytoplasmic, calcineurin dependent 2 | -0.37 | 0.01664389 | 2 |
| Tlr8 | toll-like receptor 8 | -0.36 | 0.02777452 | X |
| Asah2 | N-acylsphingosine amidohydrolase 2 | -0.35 | 0.03124079 | 19 |
| Cd93 | CD93 antigen | -0.35 | 7.86E-05 | 2 |
| Gm44250 | predicted gene, 44250 | -0.31 | 0.04195963 | 6 |
| Ulk2 | unc-51 like kinase 2 | -0.31 | 0.03138016 | 11 |
| Sft2d2 | SFT2 domain containing 2 | -0.3 | 0.00475956 | 1 |
| Irak4 | interleukin-1 receptor-associated kinase 4 | -0.29 | 0.03692125 | 15 |
| Il6ra | interleukin 6 receptor, alpha | -0.28 | 0.02813309 | 3 |
| Rab8b | RAB8B, member RAS oncogene family | -0.26 | 0.03806948 | 9 |
| Dennd5a | DENN/MADD domain containing 5A | -0.26 | 0.00588127 | 7 |
| Smad2 | small ArfGAP 2 | -0.24 | 0.03936691 | 4 |
| Dram2 | DNA-damage regulated autophagy modulator 2 | -0.23 | 0.04418227 | 3 |

Molecular mechanisms of iAs toxicity can also be inferred by our analysis given the preference of certain families of transcription factors, particularly those with zinc finger domains (i.e., ZBTB or zf-C2H2/GATA families) being perturbed after iAs exposure (**Table 2-16**). Indeed, there is evidence that iAs interacts with zinc finger motifs on DNA repair proteins, a mechanism that can partially explain arsenic’s synergistic toxic effects with certain genotoxic environmental exposures such as benzo[a]pyrene and radiation (Qing 2008; Wang 2013; Zhou 2011; and reviewed in Zhou 2021) Further, zinc homeostasis is crucial for regulating zinc-finger transcription factors (since zinc is required to form the domain that binds to DNA) (Choi 2014; Duffy 2004) and macrophage pro-inflammatory function (Gao 2018). Other results of our analysis also support that arsenic plays a role in disrupting calcium homeostasis (Florea 2005), and one study using a non-mammalian (catfish) model has demonstrated this in macrophages (Banerjee 2011). For example, the Fantom transcriptional network generated for the iAs treated female-derived M1 downregulated group shows regulation of several calcium relevant proteins, including Gpr18 (a G protein-coupled receptor that regulates Rho signal transduction (Console-Bram 2014; Sah 2000)), TUSC2 (a mitochondrial calcium regulator (Uzhachenko 2014)), and Nucb2 (which has suggested roles in calcium level maintenance (Skorupska 2020) (**Fig. 2-5C**). These findings suggest that arsenic disrupts ion homeostasis, and this should be considered in future studies focused on elucidating iAs toxicity as well as those focusing on the health effects of metal mixtures.

**Table 2-8.**
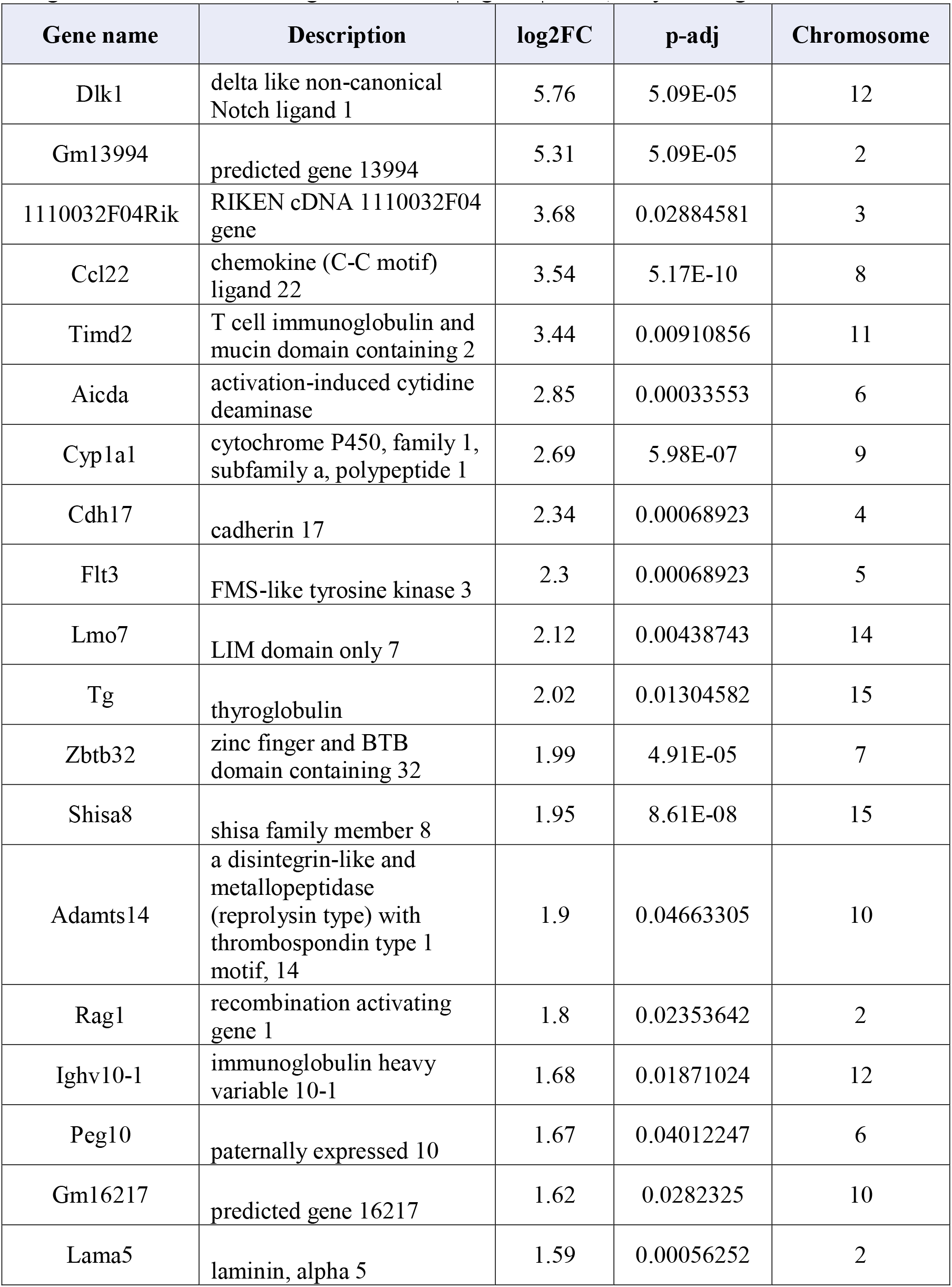
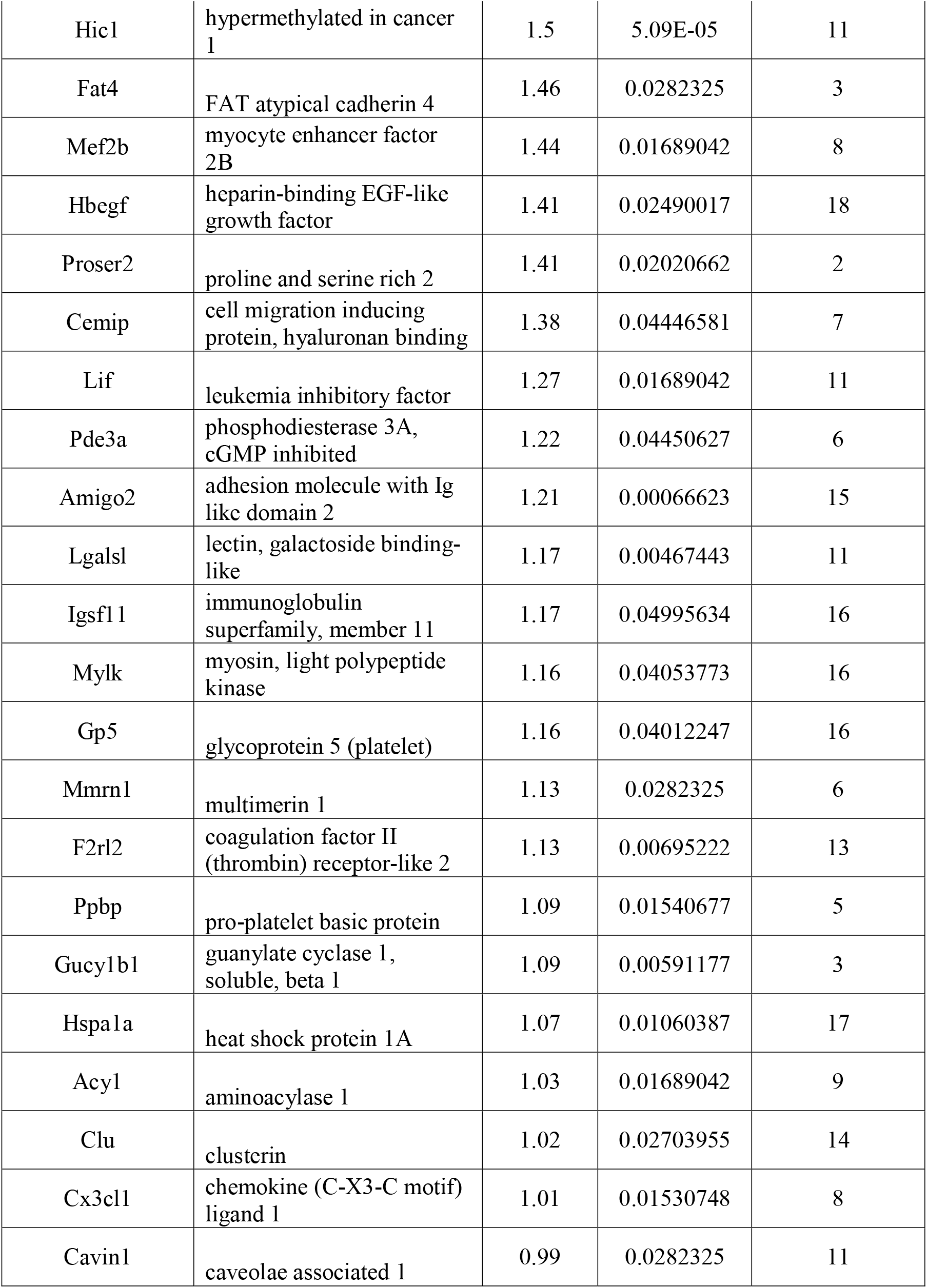
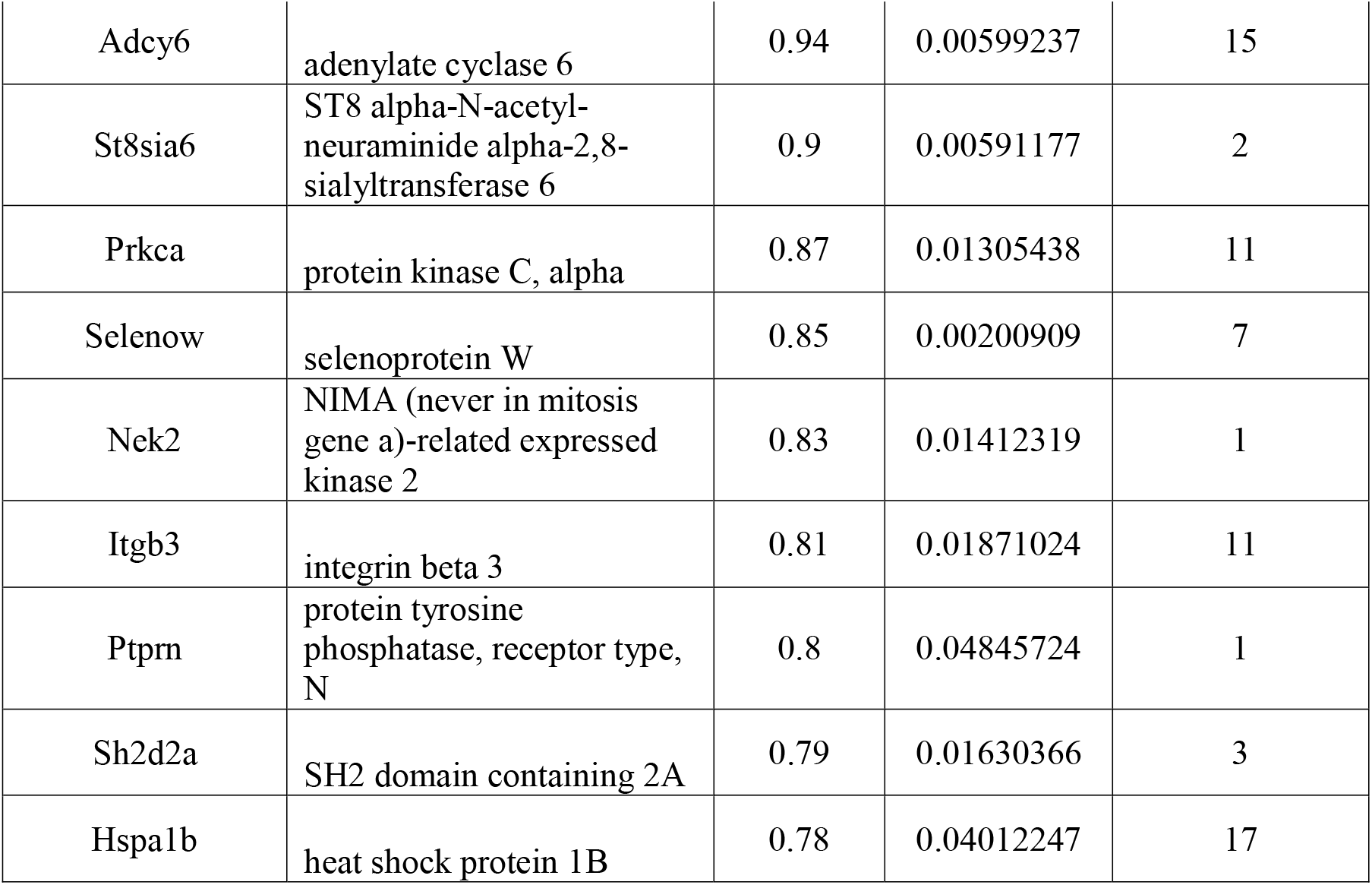
Top 50 upregulated genes in iAs-treated M0 male-derived BMDMs. If fewer than 50 genes met the more stringent cutoff of |log2FC| > 0.2, only those genes are listed.

**Table 2-9.** Top 50 downregulated genes in iAs-treated M0 male-derived BMDMs. If fewer than 50 genes met the more stringent cutoff of |log2FC| > 0.2, only those genes are listed.

| Gene name | Description | log2FC | p-adj | Chromosome |
| --- | --- | --- | --- | --- |
| Ccr2 | chemokine (C-C motif) receptor 2 | -1.32 | 1.98E-10 | 9 |
| Fcrls | Fc receptor-like S, scavenger receptor | -1.26 | 8.59E-05 | 3 |
| Cx3cr1 | chemokine (C-X3-C motif) receptor 1 | -1.04 | 0.00513786 | 9 |
| Rnd3 | Rho family GTPase 3 | -1 | 0.00467443 | 2 |
| Hpgd | hydroxyprostaglandin dehydrogenase 15 (NAD) | -0.96 | 0.00929166 | 8 |
| Mcam | melanoma cell adhesion molecule | -0.82 | 0.04012247 | 9 |
| Camp | cathelicidin antimicrobial peptide | -0.81 | 0.00279441 | 9 |
| Pglyrp1 | peptidoglycan recognition protein 1 | -0.69 | 0.01758799 | 7 |
| Slc9a9 | solute carrier family 9 (sodium/hydrogen exchanger), member 9 | -0.56 | 0.00014377 | 9 |
| Lpar6 | lysophosphatidic acid receptor 6 | -0.49 | 0.00694006 | 14 |
| Gas6 | growth arrest specific 6 | -0.48 | 0.00676698 | 8 |
| Gas7 | growth arrest specific 7 | -0.38 | 0.01996044 | 11 |
| Otulinl | OTU deubiquitinase with linear linkage specificity like | -0.36 | 0.04012247 | 15 |
| Wwp1 | WW domain containing E3 ubiquitin protein ligase 1 | -0.31 | 0.00438743 | 4 |

**Table 2-10.**
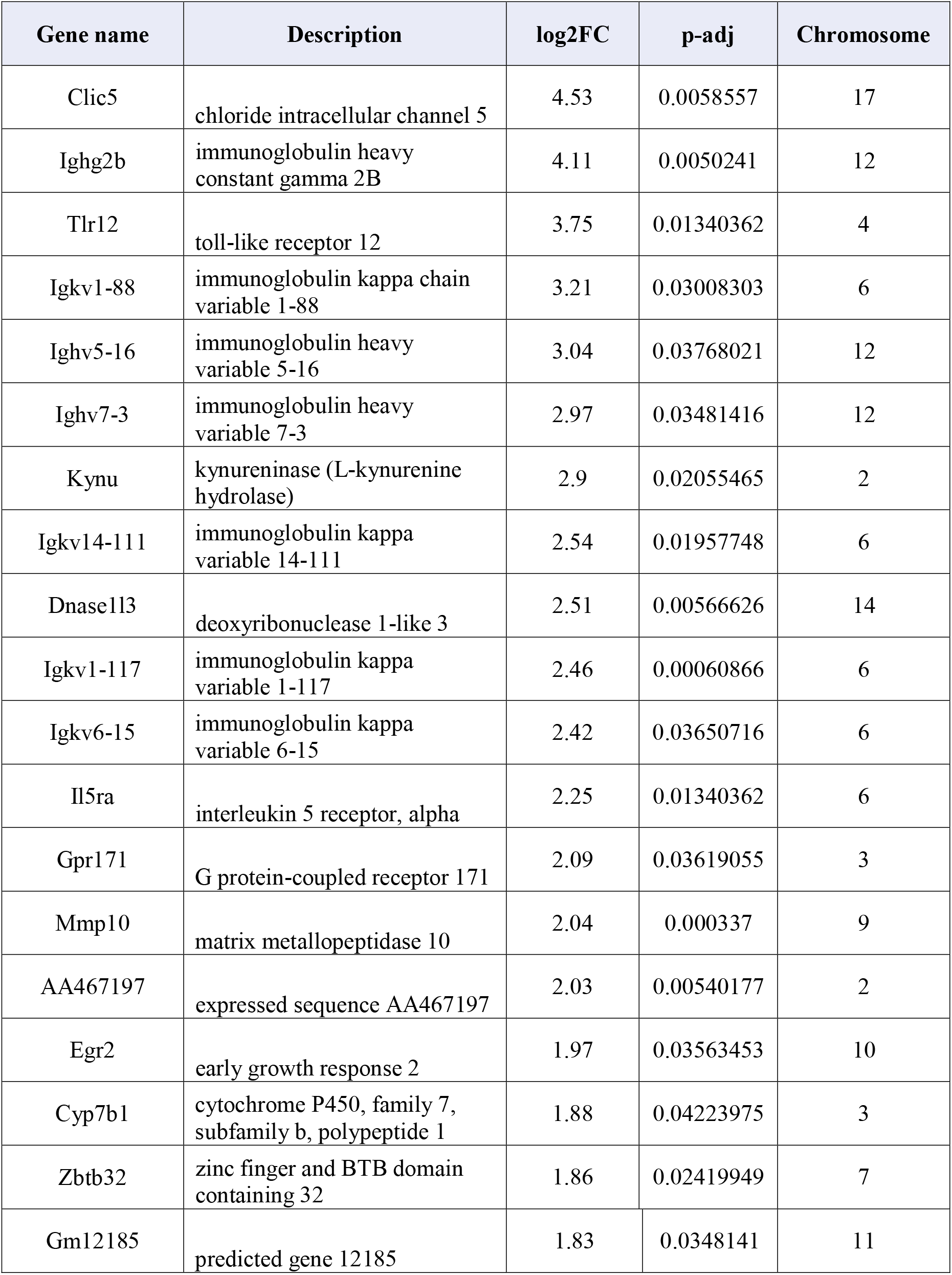
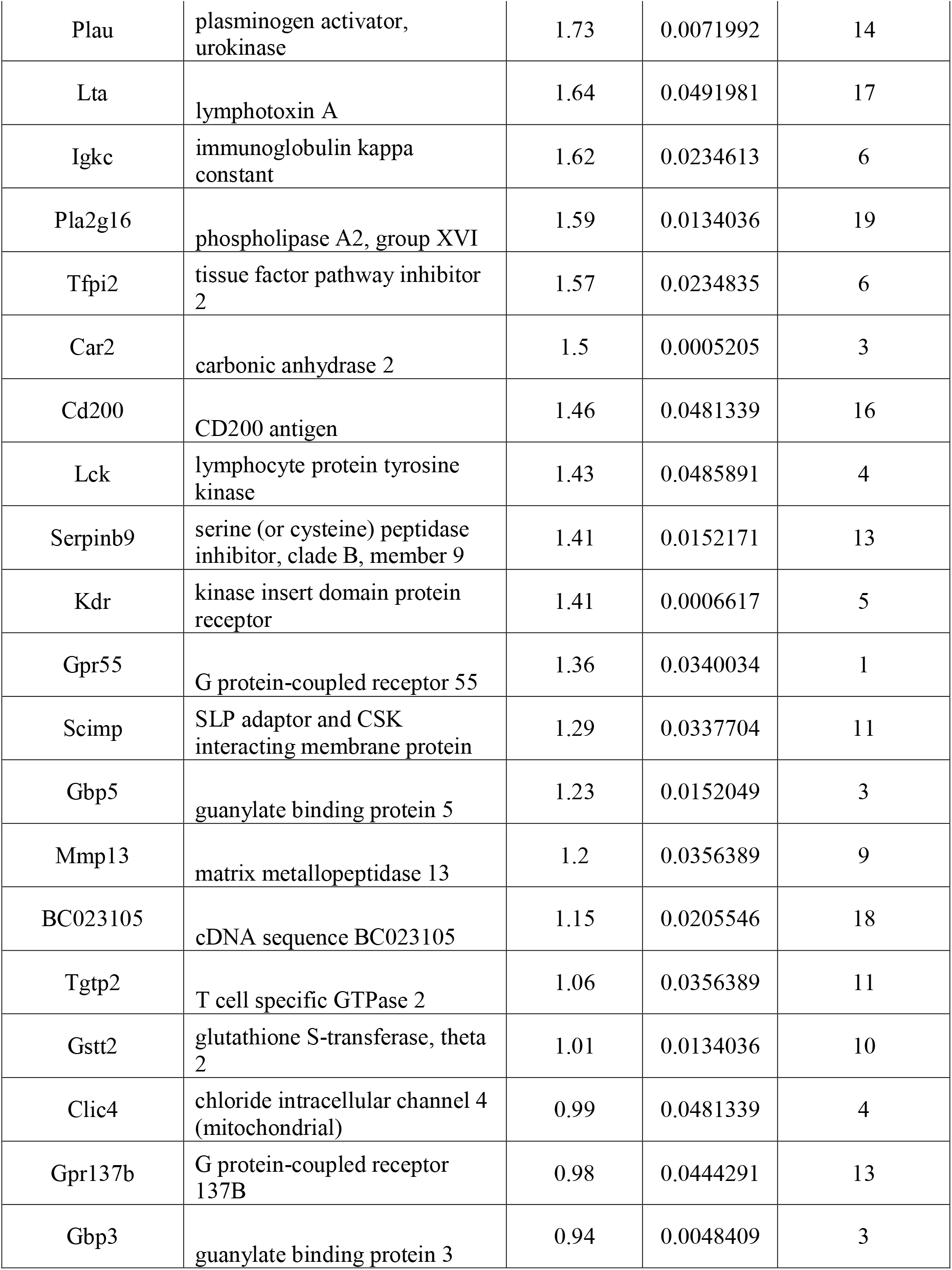
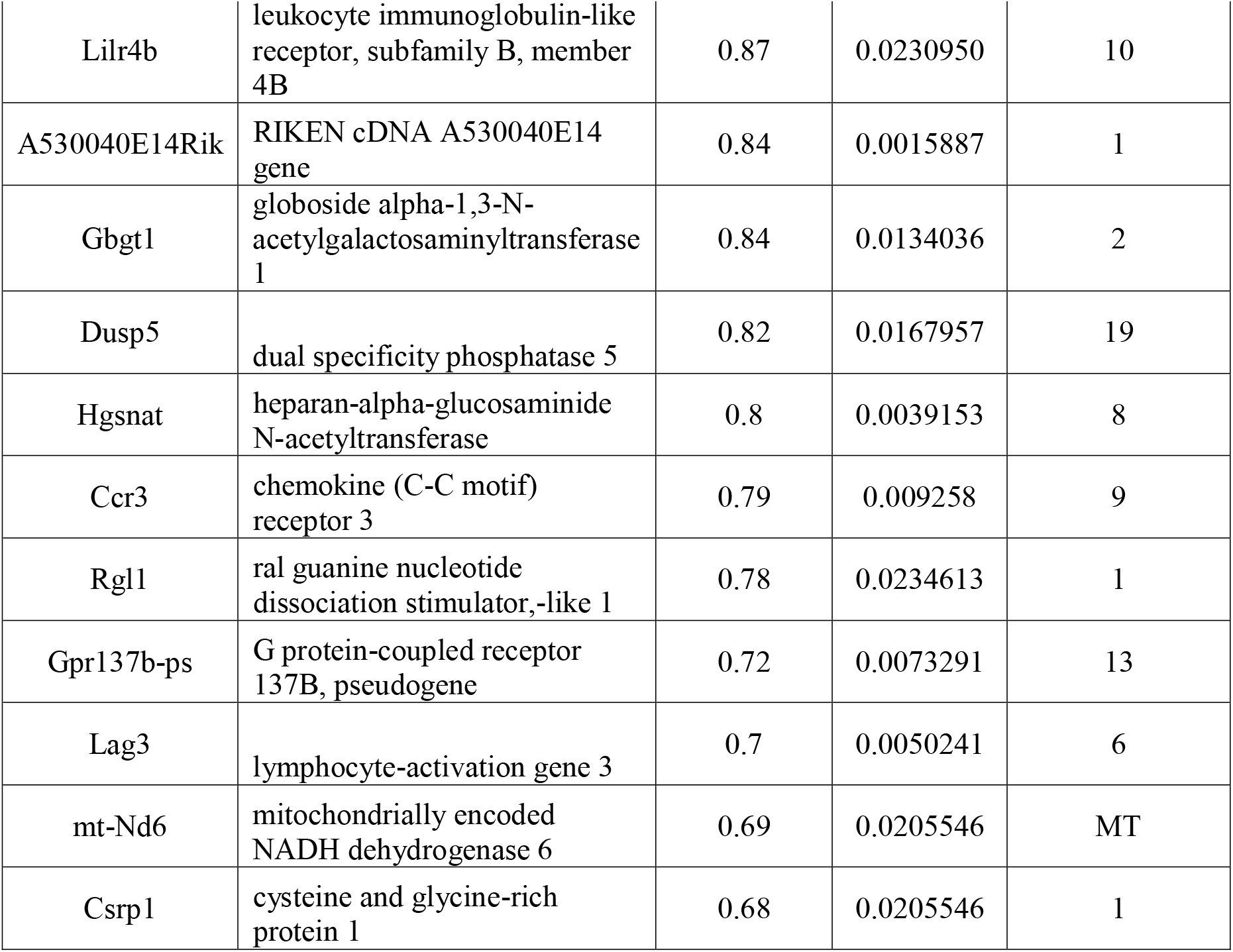
Top 50 upregulated genes in iAs-treated M1 male-derived BMDMs. If fewer than 50 genes met the more stringent cutoff of |log2FC| > 0.2, only those genes are listed.

**Table 2-11.**
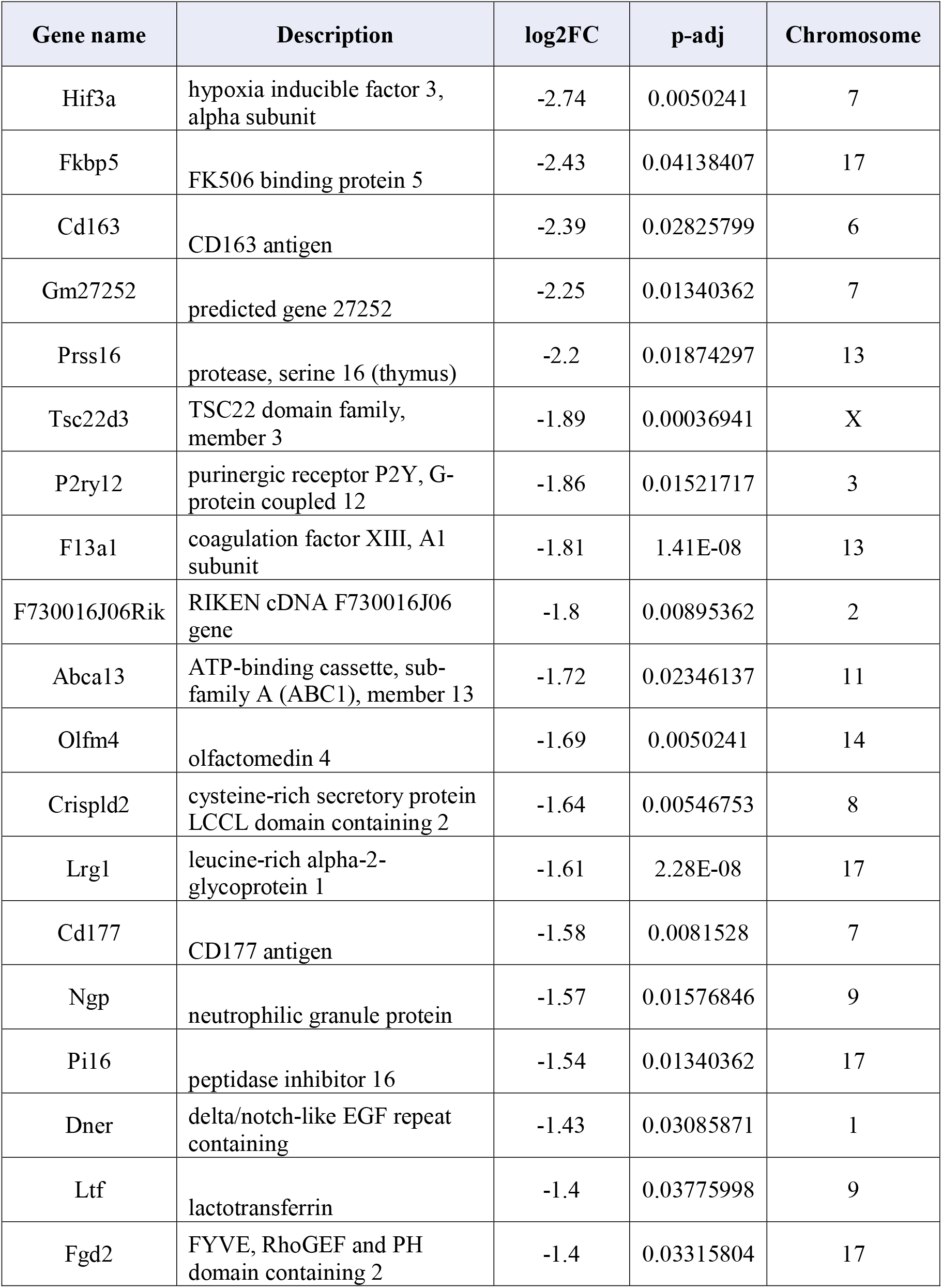
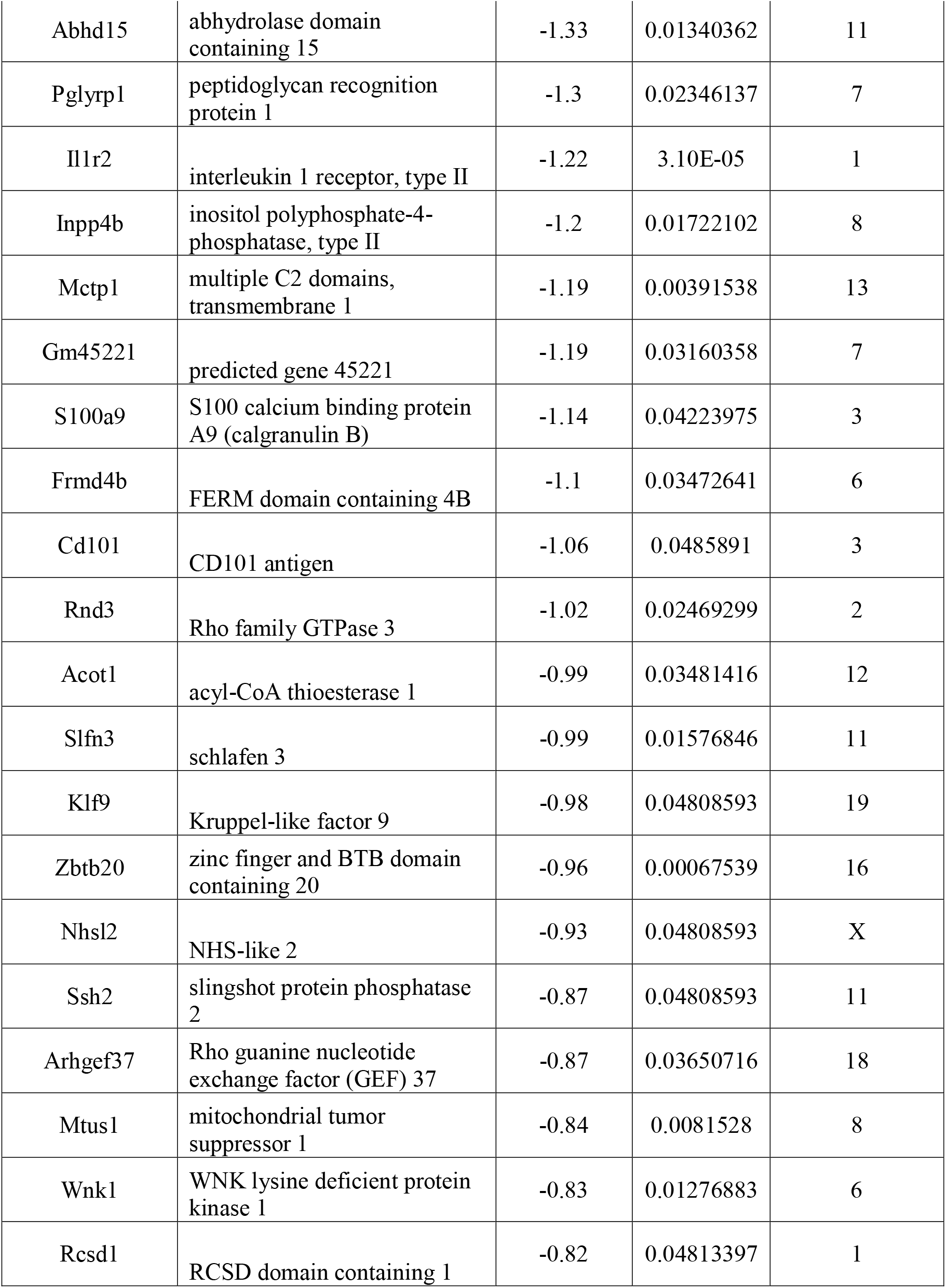
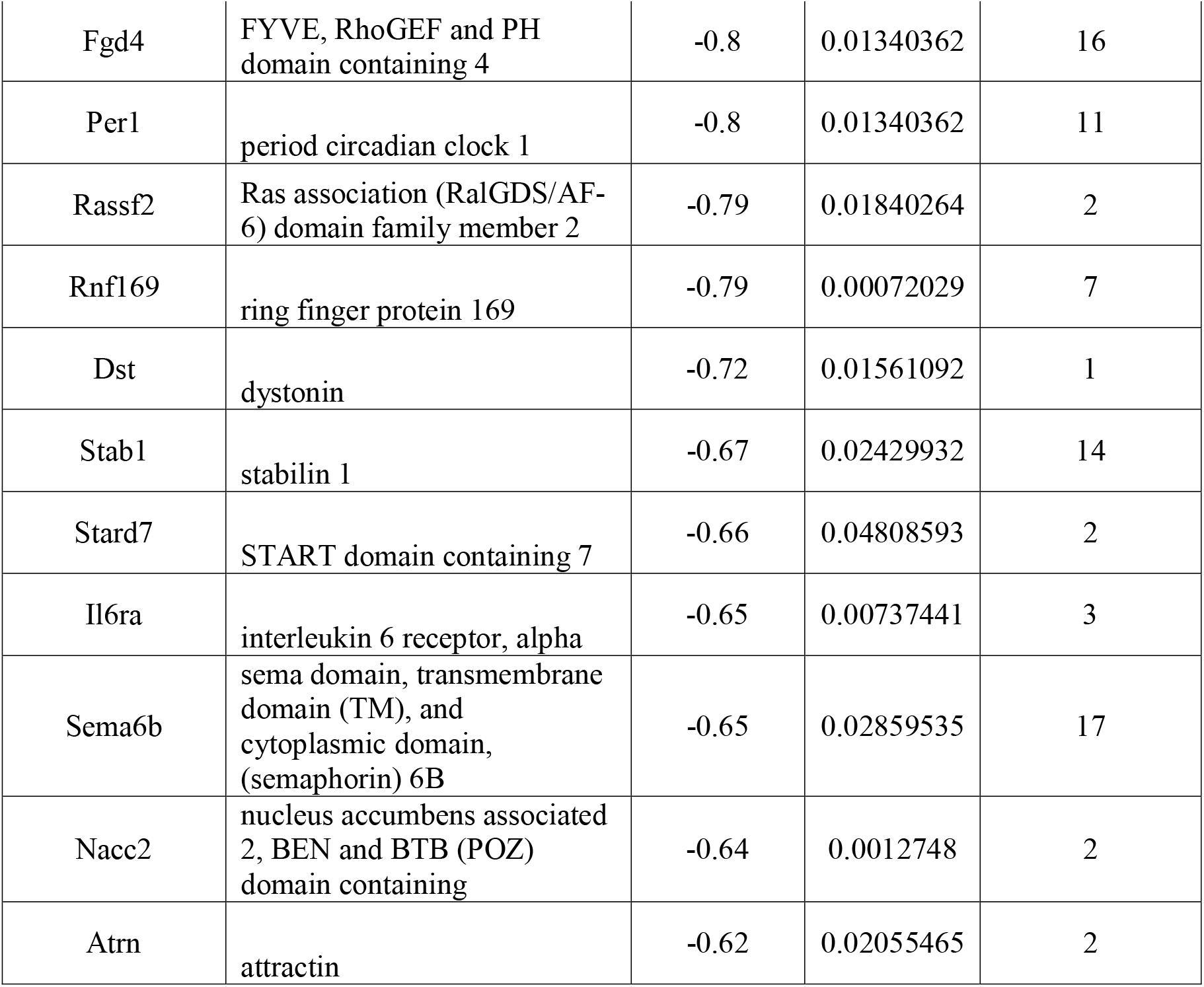
Top 50 downregulated genes in iAs-treated M1 male-derived BMDMs. If fewer than 50 genes met the more stringent cutoff of |log2FC| > 0.2, only those genes are listed.

**Table 2-12.** Top 50 upregulated genes in iAs-treated M2 male-derived BMDMs. If fewer than 50 genes met the more stringent cutoff of |log2FC| > 0.2, only those genes are listed.

| Gene name | Description | log2FC | p-adj | Chromosome |
| --- | --- | --- | --- | --- |
| Calca | calcitonin/calcitonin-related polypeptide, alpha | 1.93 | 0.00160036 | 7 |
| Spry1 | sprouty RTK signaling antagonist 1 | 1.53 | 0.0375221 | 3 |
| Serpina3f | serine (or cysteine) peptidase inhibitor, clade A, member 3F | 1.32 | 0.01736253 | 12 |
| Kcnj5 | potassium inwardly-rectifying channel, subfamily J, member 5 | 1.25 | 0.01101952 | 9 |
| Serpina3g | serine (or cysteine) peptidase inhibitor, clade A, member 3G | 1.2 | 1.31E-05 | 12 |
| Prelp | proline arginine-rich end leucine-rich repeat | 1.18 | 0.00915911 | 1 |
| Ltbp1 | latent transforming growth factor beta binding protein 1 | 1.14 | 6.18E-06 | 17 |
| Itga2b | integrin alpha 2b | 1.11 | 0.01275565 | 11 |
| Gpx3 | glutathione peroxidase 3 | 1.09 | 2.83E-06 | 11 |
| Fkbp5 | FK506 binding protein 5 | 1.09 | 0.02417942 | 17 |
| Clu | clusterin | 1.08 | 0.00946444 | 14 |
| Ctla2a | cytotoxic T lymphocyte-associated protein 2 alpha | 1.07 | 0.01736253 | 13 |
| Samd14 | sterile alpha motif domain containing 14 | 1.06 | 0.0084944 | 11 |
| Mmrn1 | multimerin 1 | 1.05 | 0.0476662 | 6 |
| Kbtbd11 | kelch repeat and BTB (POZ) domain containing 11 | 1.04 | 0.03421428 | 8 |
| Angpt1 | angiopoietin 1 | 0.99 | 0.02593944 | 15 |
| Smo | smoothened, frizzled class receptor | 0.99 | 0.03636506 | 6 |
| Serpine2 | serine (or cysteine) peptidase inhibitor, clade E, member 2 | 0.97 | 6.24E-05 | 1 |
| CT010467.1 | 18s RNA, related sequence 5 | 0.96 | 0.02701945 | 17 |
| Nrgn | neurogranin | 0.95 | 0.04833115 | 9 |
| Plxna4 | plexin A4 | 0.94 | 0.02975538 | 6 |
| Pde5a | phosphodiesterase 5A,<br>cGMP-specific | 0.92 | 0.0498527 | 3 |
| Lgalsl | lectin, galactoside binding-like | 0.91 | 0.02902505 | 11 |
| Adcy6 | adenylate cyclase 6 | 0.89 | 0.02391291 | 15 |
| Myo1d | myosin ID | 0.86 | 0.01781308 | 11 |
| Cd34 | CD34 antigen | 0.78 | 0.02391291 | 1 |
| Il1rl1 | interleukin 1 receptor-like 1 | 0.71 | 0.02701945 | 1 |
| Dhcr24 | 24-dehydrocholesterol<br>reductase | 0.67 | 0.04833115 | 4 |
| Scd2 | stearoyl-Coenzyme A<br>desaturase 2 | 0.66 | 0.01736253 | 19 |
| F5 | coagulation factor V | 0.64 | 0.00901269 | 1 |
| Pmepa1 | prostate transmembrane<br>protein, androgen induced 1 | 0.6 | 0.0375221 | 2 |
| Cd81 | CD81 antigen | 0.57 | 0.00543873 | 7 |
| Nedd4 | neural precursor cell<br>expressed, developmentally<br>down-regulated 4 | 0.46 | 0.0476662 | 9 |
| Vcam1 | vascular cell adhesion<br>molecule 1 | 0.44 | 0.00086233 | 3 |
| AI506816 | expressed sequence<br>AI506816 | 0.42 | 0.01015899 | 5 |

**Table 2-13.** Top 50 downregulated genes in iAs-treated M2 male-derived BMDMs. If fewer than 50 genes met the more stringent cutoff of |log2FC| > 0.2, only those genes are listed.

| Gene name | Description | log2FC | p-adj | Chromosome |
| --- | --- | --- | --- | --- |
| Ccl24 | chemokine (C-C motif)<br>ligand 24 | -1.17 | 0.02391291 | 5 |
| Fcrls | Fc receptor-like S, scavenger<br>receptor | -1.04 | 0.00019749 | 3 |
| Penk | preproenkephalin | -1.02 | 0.00296886 | 4 |
| Sirpb1c | signal-regulatory protein beta<br>1C | -1 | 0.03188178 | 3 |
| Ccr2 | chemokine (C-C motif)<br>receptor 2 | -0.9 | 0.02902505 | 9 |
| Arl5c | ADP-ribosylation factor-like<br>5C | -0.72 | 0.00210925 | 11 |
| Slc16a3 | solute carrier family 16<br>(monocarboxylic acid<br>transporters), member 3 | -0.63 | 0.02902505 | 11 |
| Gpr132 | G protein-coupled receptor<br>132 | -0.56 | 0.03188178 | 12 |
| Tbc1d8 | TBC1 domain family,<br>member 8 | -0.39 | 0.02701945 | 1 |
| Sema4a | sema domain,<br>immunoglobulin domain (Ig),<br>transmembrane domain (TM)<br>and short cytoplasmic<br>domain, (semaphorin) 4A | -0.38 | 0.01275565 | 3 |
| Lpcat2 | lysophosphatidylcholine<br>acyltransferase 2 | -0.38 | 0.03032979 | 8 |

**Table 2-14.** STRING network characteristics. If no data is shown for a group, no network was generated given our input settings.

| Sex | Stimulation | FC direction | # of nodes | # of edges (actual, expected) | Avg. node degree | Avg. clustering coefficient | PPI enrichment p-value |
| --- | --- | --- | --- | --- | --- | --- | --- |
| Female | M0 | Down | 25 | 1, 1 | 0.08 | 0.08 | 0.454 |
|  |  | Up | 134 | 60, 32 | 0.896 | 0.274 | 5.7 x 10<sup>-6</sup> |
|  | M1 | Down | 301 | 294, 193 | 1.95 | 0.321 | 8.96 x 10<sup>-12</sup> |
|  |  | Up | 298 | 225, 116 | 1.51 | 0.282 | < 1 x 10<sup>-16</sup> |
|  | M2 | Up | 65 | 12, 6 | 0.369 | 0.2 | 0.0321 |
| Male | M0 | Up | 61 | 10, 7 | 0.328 | 0.164 | 0.148 |
|  | M1 | Down | 67 | 1, 6 | 0.0299 | 0.0299 | 0.997 |
|  |  | Up | 42 | 3, 3 | 0.143 | 0.143 | 0.626 |
|  | M2 | Up | 33 | 5, 1 | 0.303 | 0.242 | 0.00313 |

To highlight immunologically relevant genes in our STRING networks, we utilized Cytoscape. Briefly, tables from STRING interaction networks were uploaded to Cytoscape and nodes that were immunologically relevant were color-coded based on categorical function. A total of 8 categories are highlighted: integrin signaling (green), coagulation factors (red), chemotaxis and chemokine signaling (orange), pattern recognition and antigen presentation (pink), splicing machinery (yellow), lipid metabolism (purple), complement system (blue), and cytokine signaling (aqua) (**Fig. 2-4**).

Genes encoding proteins related to integrin signaling (e.g., *Itgab3*, *Itga2*, *Itgax*) were upregulated in iAs-treated Female M0 and M1 groups (**Figs. 2-4A&D**), as well as the iAs-treated Male M0 group (**Fig. 2-4C**). Genes encoding proteins involved in coagulation processes (e.g., *F2rl2*, *F2r*, *F2rl3*, *Mmrn1*, *F5*) were upregulated in the iAs treated Female M0 group (**Fig. 2-4A**), the iAs-treated Female M1 group (**Fig. 2-4B**), and iAs-treated Female and Male M2 groups (**Figs. 2-4H&I**). Protein-coding genes involved in chemotaxis and chemokine signaling (e.g., *Ppbp*, *Ccl22*, *Cx3cl1*, *Ccr6*, *Lif*, *Hbegf*) were upregulated in the iAs-treated Female and Male M0 groups (**Figs. 2-4A&C**), while others (*Mmps*) were upregulated in the iAs-exposed male-derived M1 and female-derived M2 group (**Figs. 2-4F&H**). Pattern recognition and antigen presentation genes (e.g., *Ctla4*, *H2-m2*) were upregulated in iAs-treated female-derived groups across all 3 stimulations (**Figs. 2-4A, D, and H**), while others (e.g., *H2-d1*, *Cd28*, *Tlr7*, *Tlr8*, *Myd88*) were downregulated in iAs-treated Female M0 and M1 groups (**Figs. 2-4B&E**). Genes related to splicing machinery (e.g., *Srsfs* and *Nxf1*) were upregulated in the iAs-treated Female M1 group (**Fig. 2-4D**). Protein-coding genes controlling lipid metabolism (e.g., *Pla2g16*, *Alox12*, *Lpl*, *Fabp3*, *Acoxl*) were also upregulated in the iAs-treated Female M1 group (Fig. 3D). Protein-coding genes involved in the complement system (e.g., *C1qa*, *C1qb*, *C1qc*, *C3*, *C5ar1*) were downregulated in the iAs-treated Female M1 group (**Fig. 2-4E**). Finally, certain protein-coding genes involved in cytokine signaling (e.g., *Flt3*, *Fgfr1*, *IL5ra*, *Egr1*, *IL12a*, *Csf1*, *Tgfb1i1*) were upregulated in iAs-treated Females across the three stimulations (**Figs. 2-4A, D, and H**), while others (e.g., *Il6ra*, *Irf8*, *Ifitm2*, *Ifitm3*, *Ifngr1*, *Ifnar2*, *Csf1r*, *Tgfb1*) were downregulated in the iAs-treated Female M1 group only (**Fig. 2-4E**).

### iAs-induced Changes in Non-protein-coding Transcripts

It is controversial as to what extent non-protein coding transcripts, such as small RNAs that regulate gene expression or pseudogenes with no predicted function, are biologically relevant (Flores-Concha 2020; Palazzo 2020). At the same time, several non protein-coding transcripts have received attention for their relevance in regulating the immune response (Flores-Concha 2020). In addition to protein-coding genes, our RNA-seq data set revealed iAs-induced changes in expression (52 upregulated vs. only 12 downregulated total across our DEG lists) of antisense genes, pseudogenes and transcripts, genes to be experimentally confirmed (TECs), as well as non-coding RNAs (ncRNAs) including long intergenic non-coding RNAs (lincRNAs), microRNAs (miRNAs), and ribosomal RNAs (rRNAs) in male and female BMDMs across stimulation groups (**Table 2-15**).

**Table 2-15.**
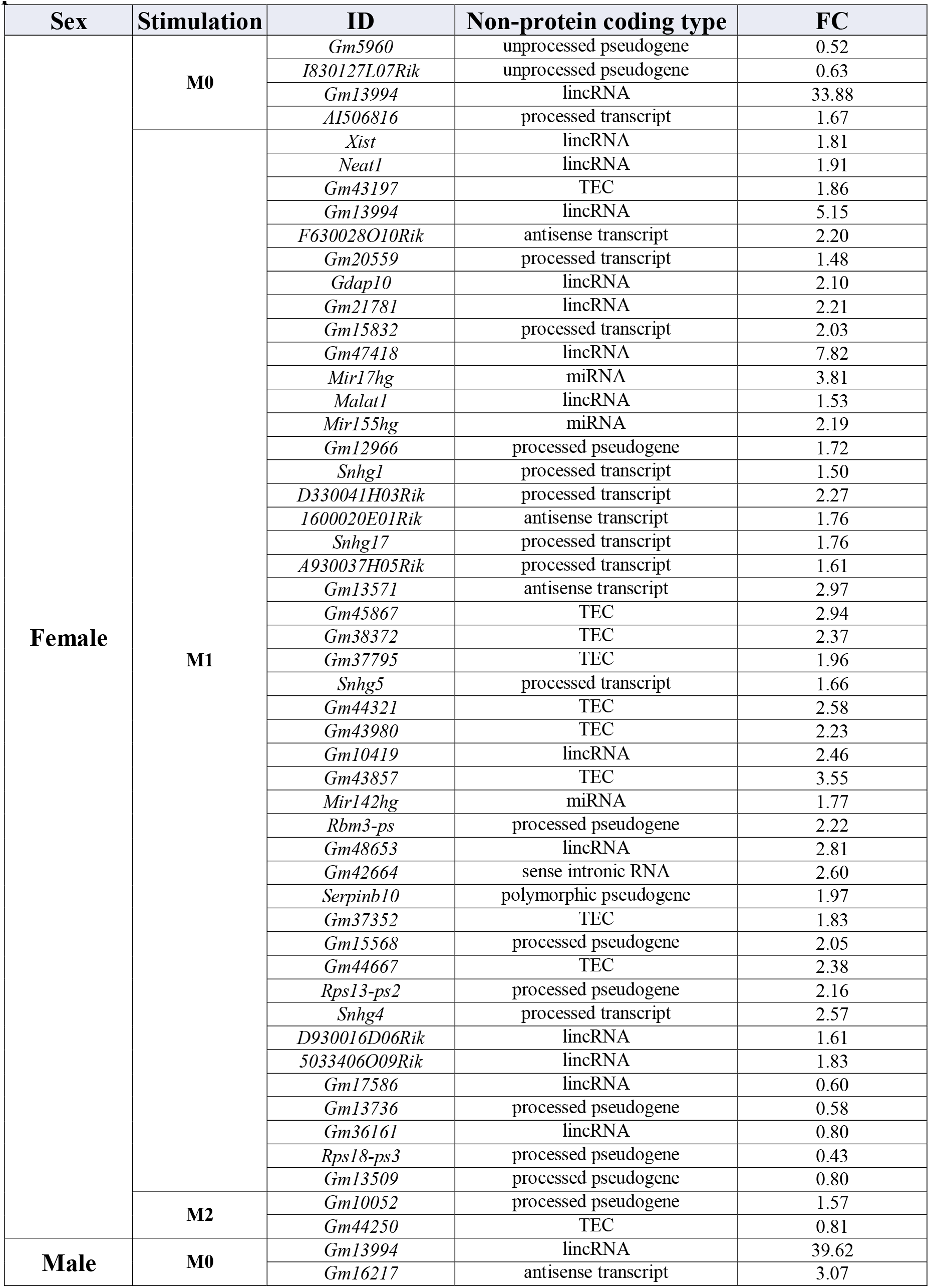
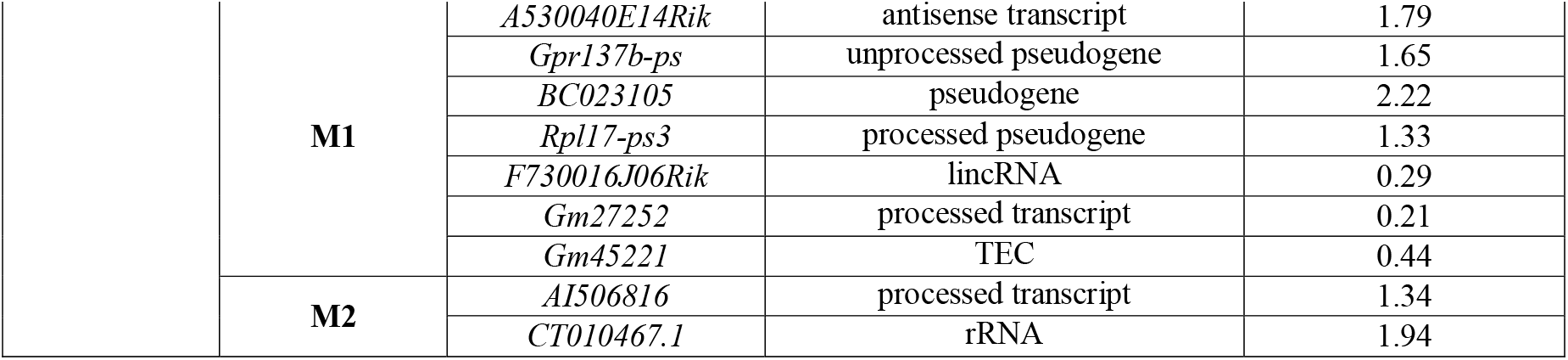
Non-protein coding transcripts differentially expressed in female and male iAs exposed BMDMs.

### Differential Expression of Transcription Factors in BMDMs in a Sex- and Stimulation-Dependent Manner

Transcription factors maintain transcriptomic homeostasis and regulate global responses to external stimuli (Vaquerizas 2009; Wilkinsin 2017). Because of their broad influence in regulating transcription of downstream genes and relevance in disease, we identified transcription factors perturbed by iAs exposure within each sex and across stimulation groups in our DEG lists. These transcription factors, along with their categorized transcription factor family and fold change values, are highlighted in **Table 2-16**. Of the 48 total transcription factors identified across our comparison groups, 23 (ZBTB or zf-C2H2/GATA families) have zinc finger-containing domains. Additional families of transcription factors that were perturbed by arsenic in our DEG lists included the MYB family (4 total, all upregulated in the female-derived M1 group), the HMG/HMGI/HMGY family (3 total, 2 upregulated in the iAs-treated female-derived M1 group and 1 upregulated in the iAs-treated male-derived M0 group), the bHLH family (3 total, 1 upregulated in iAs-treated female-derived M0 group, 1 upregulated in the iAs treated female-derived M1 group, and 1 downregulated in the iAs-treated female-derived M1 group), the SRF family (3 total, 1 upregulated in iAs-treated the female-derived M0 group, 1 upregulated in the iAs-treated male-derived M0 group, and 1 downregulated in the iAs-treated female-derived M1 group), and the IRF family (2 total, 1 downregulated in the iAs-treated female-derived M0 group and 1 downregulated in the iAs-treated female-derived M1 group).

**Table 2-16.**
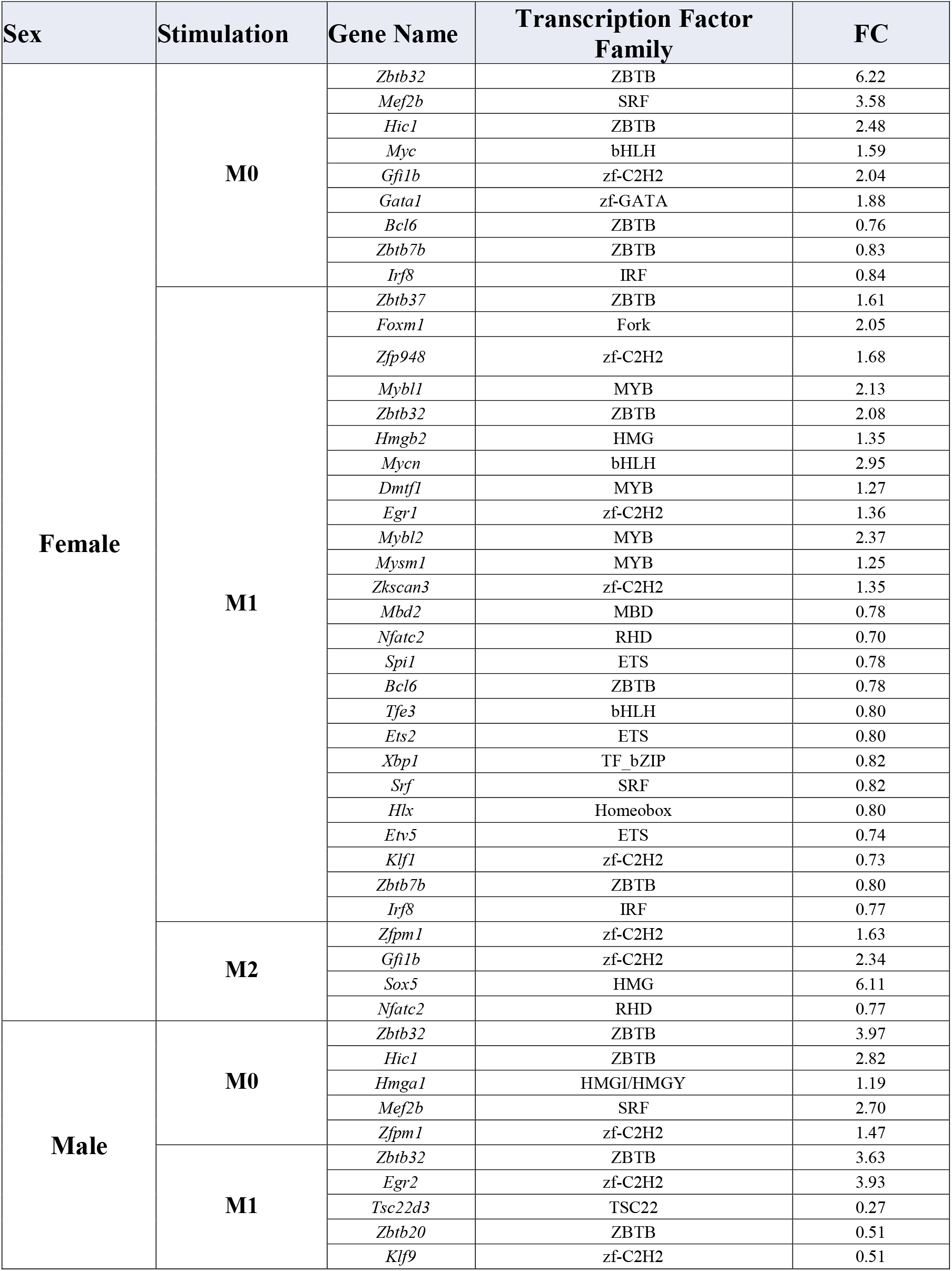
Transcription factors perturbed by arsenic exposure in female and male BMDMs across stimulation groups. None were significant in the Male M2 group.

### FANTOM Regulatory Interactions

FANTOM4 EdgeExpressDB (Ravasi 2010; Severin 2009) was used to visualize regulatory interactions and transcriptional networks for each of our DEG lists (with cutoff thresholds applied as previously described). Only significant networks were generated for the iAs-treated Female M0 upregulated group, the iAs-treated Female M1 up- and downregulated groups, and the iAs-treated Female M2 upregulated group (**Fig. 2-5**). The iAs-treated female-derived M0 upregulated transcriptional network shows several genes regulated by MYBl2 and MYC (**Fig. 2-5A**). The iAs-treated female-derived M1 upregulated gene list generated networks majorly regulated by transcription factors BRCA1 and TMF1 (**Fig. 2-5B**), while the iAs-treated female-derived M1 downregulated list generated the most extensive network, with transcription factors IRF8, SP1, and ETS2 regulating expression of over 100 downstream genes (**Fig. 2-5C**). The iAs-treated female-derived M2 upregulated gene list generated a network of 7 downstream targets regulated by UHRF1 (**Fig. 2-5D**). The subnetwork views of these networks are color coded and scaled as previously described (Severin 2009); briefly, the edge color represents the source of the interaction where interactions from ChIP data are shown in green, protein-protein interactions are shown in purple, and published protein-DNA interactions are shown in yellow. The diameter and purple color of each node are scaled to the dynamics of the gene as determined by Illumina expression intensity and time course data (darker purple= higher intensity expression, wider diameter= more dynamic expression over a time course). We focus our discussion of these Fantom networks on major transcriptional regulators and highly expressed and dynamic genes.

**Figure 2-5.**
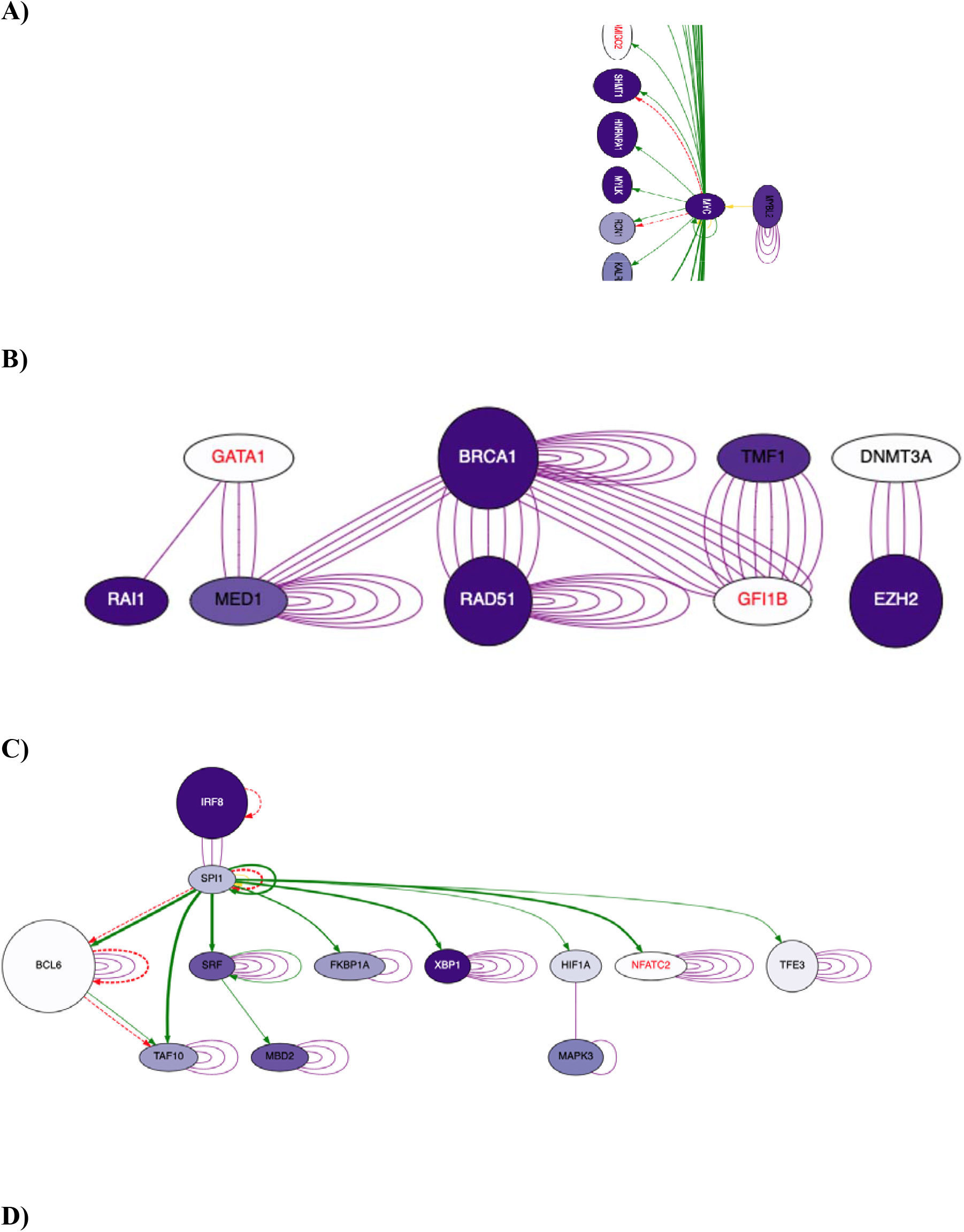
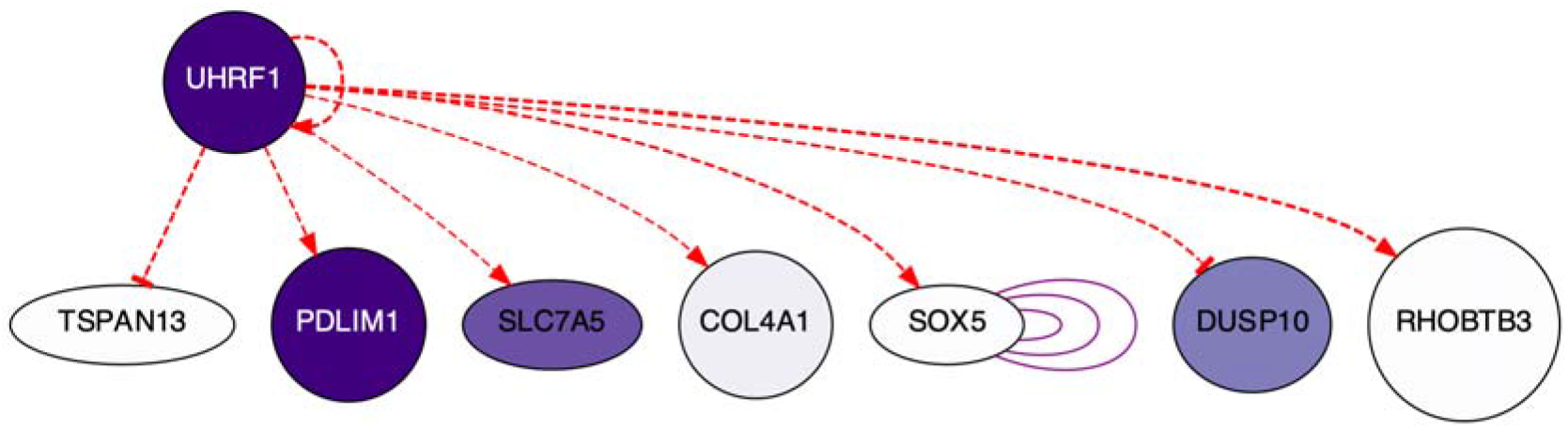
Transcriptional regulatory networks generated in Fantom for up- and downregulated genes from female and male iAs-exposed BMDMs across stimulation groups. Fantom networks were generated for DEG lists from female-derived M0 upregulated group (A), the female-derived M1 up- (B) and downregulated (C) groups, and the female-derived M2 upregulated group (D). Edge color represents the source of the interaction: ChIP data (green), protein-protein interactions (purple), published protein-DNA interactions (yellow). Multiple lines between nodes indicate multiple sources of evidence supporting the interaction and self-loops indicate evidence that a gene interacts with and/or regulates itself. Node diameter correlates with dynamic expression and purple color intensity correlates with intensity of expression. If no map was generated for a comparison group, that group is omitted from this figure.

### Differential Expression of X-linked Genes Across iAs-exposed BMDMs

Our DEG lists and subsequent analyses have revealed sex-specific changes in gene expression profiles and networks across each stimulation group. To identify genes that might be responsible for these sex-dependent findings, we highlighted X-linked genes in each of our female and male DEG lists (**Table 2-17**). The iAs-treated male stimulation groups had limited differential expression of X-linked genes, with only one having significant upregulation in the iAs-treated M0 group (*Pim2*, which was also shared with iAs-treated M0 female-derived BMDMs, and had a FC of 1.56), and only two differentially expressed X-linked genes that were unique to iAs-treated M1 males (*Tsc22d3* and *Nhsl2*, with FCs 0.27 and 0.52, respectively). iAs-treated females, on the other hand, had 24 differentially expressed X-linked genes across the three stimulations (**Table 2-17**).

**Table 2-17.**
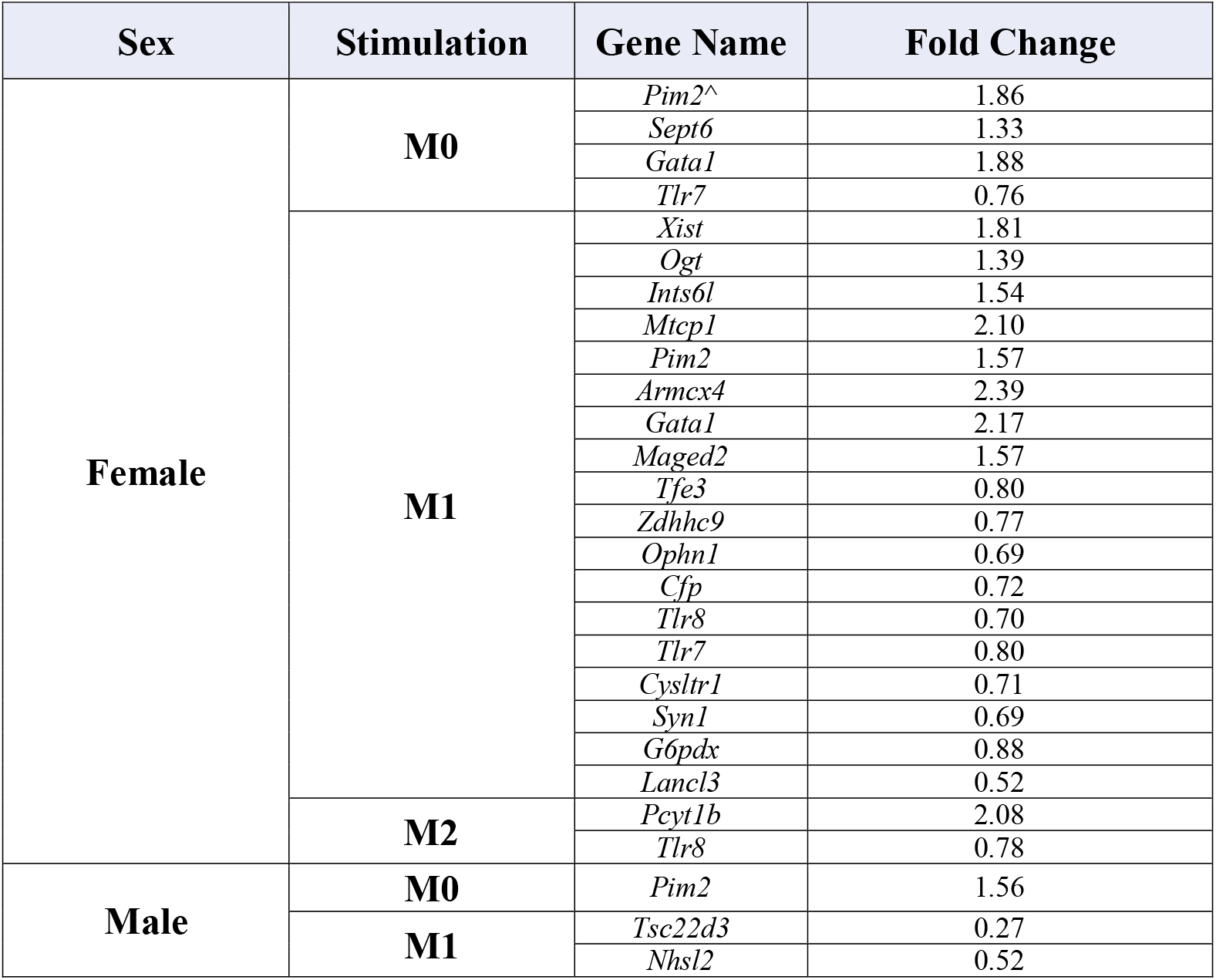
X-linked perturbed by arsenic exposure in female and male BMDMs across stimulation groups. None were significant in the Male M2 group.

### GO Enrichment Analysis of iAs-treated BMDMs

GO enrichment analysis (Ashburner 2000; GO Consortium 2021; Mi 2019) was used to identify biological processes, associated cellular components, and molecular functions that are overrepresented in up- or downregulated DEG sets from iAs-treated samples from each stimulation group across sexes. Enrichment analyses for the iAs treated female-derived M0 group showed several enriched biological processes for the upregulated gene list, including hemostasis, coagulation, wound healing, and immune system processes (**Fig. 2-6A**); enriched cellular components including the plasma membrane, perinuclear region of the cytoplasm, and the centromere (**Fig. 2-6B**); and enriched molecular functions including protein and ion binding, proteinase-activated receptor activity, and carbohydrate derivative binding (**Fig. 2-6C**). Additional immune relevant biological processes were also identified for upregulated genes from iAs-treated M0 Females (**Fig. 2-7A**) and included gens important for immune system development and response to cytokines. For the downregulated genes from iAs-treated M0 Females, we identified enriched biological processes such as regulation of the immune response and cytokine production (**Fig. 2-6D**), but no significant enrichments were found for cellular components or molecular function. For upregulated genes from iAs-treated M0 Males, we identified several enriched biological processes similar to the iAs-treated M0 Females including hemostasis and blood coagulation, but also others such as positive regulation of macromolecule metabolic processes and neutrophil and granulocyte chemotaxis (**Fig. 2-6E**); enriched cellular components including the cell surface, extracellular region, and the alphaV-beta3 integrin-PKCalpha complex (**Fig. 2-6F**); and enriched molecular functions including signaling receptor binding, integrin binding and misfolded protein binding (**Fig. 2-6G**). Additional immune-relevant significantly enriched biological processes, including positive regulation of leukocyte differentiation, leukocyte chemotaxis, and response to wounding, were also identified for the upregulated iAs-treated M0 male-derived gene list (**Fig. 2-7B**). No significant enrichments for biological processes, cellular compartments, or molecular function were found for the downregulated iAs-treated M0 male-derived gene list.

**Figure 2-6.**
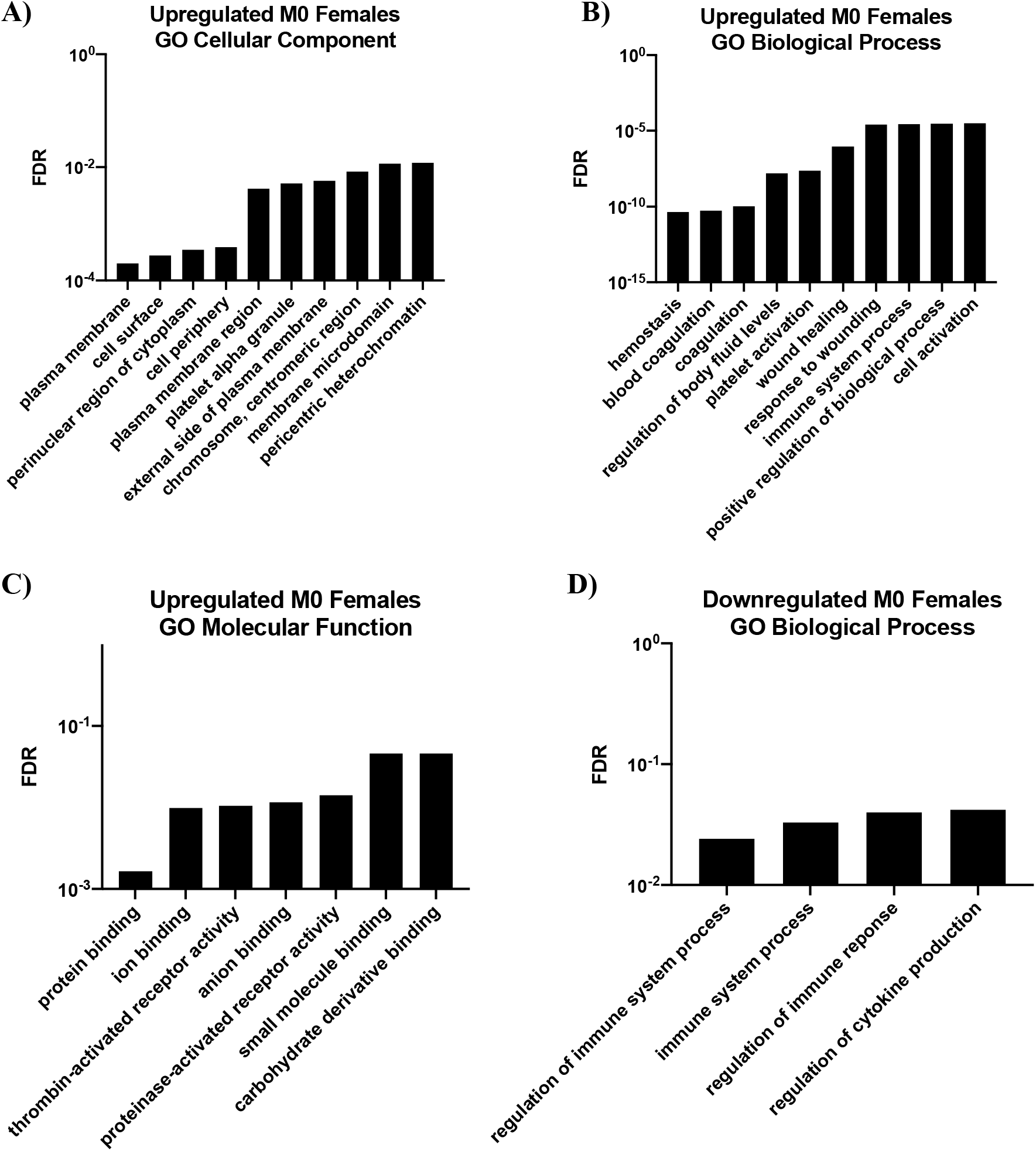
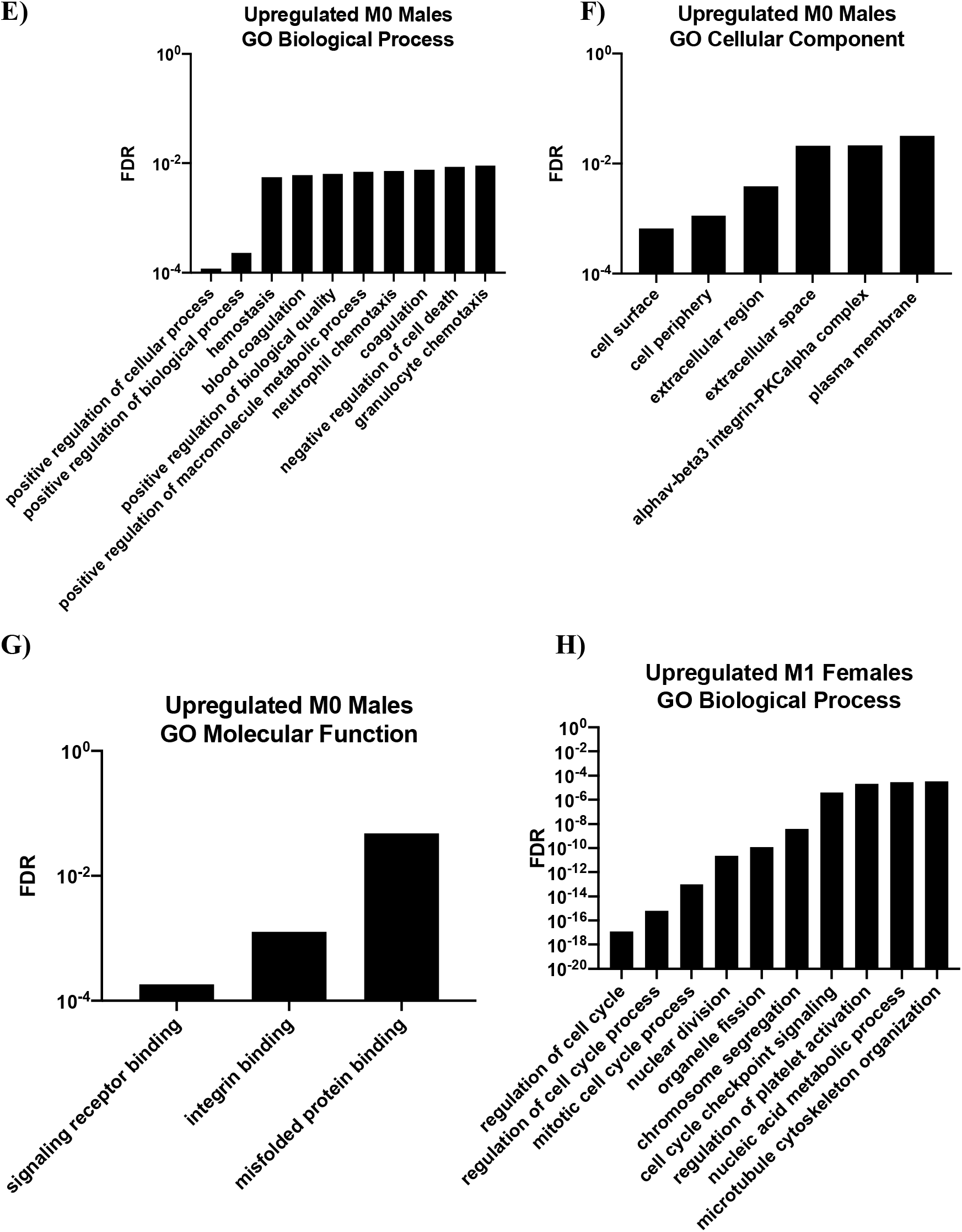
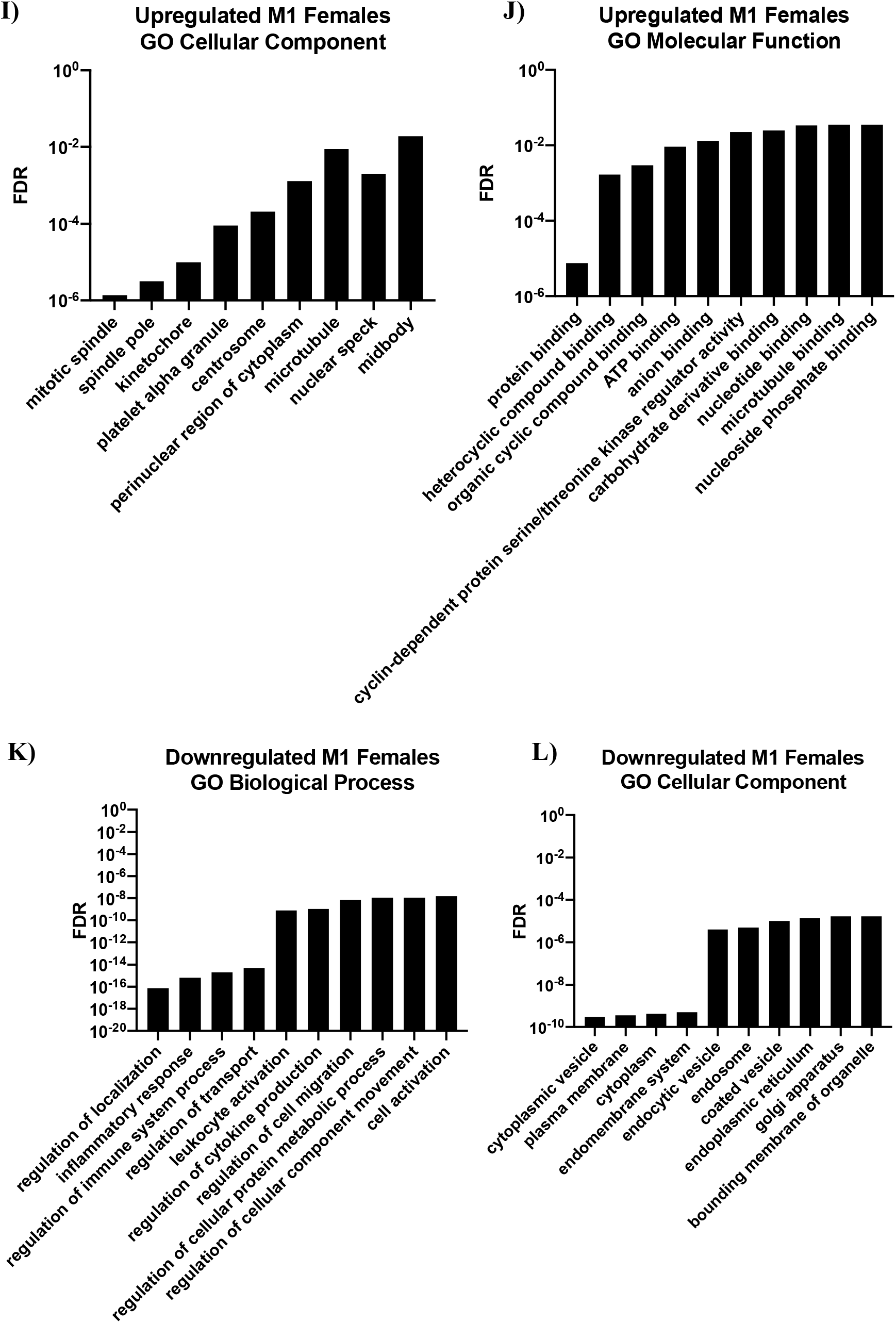
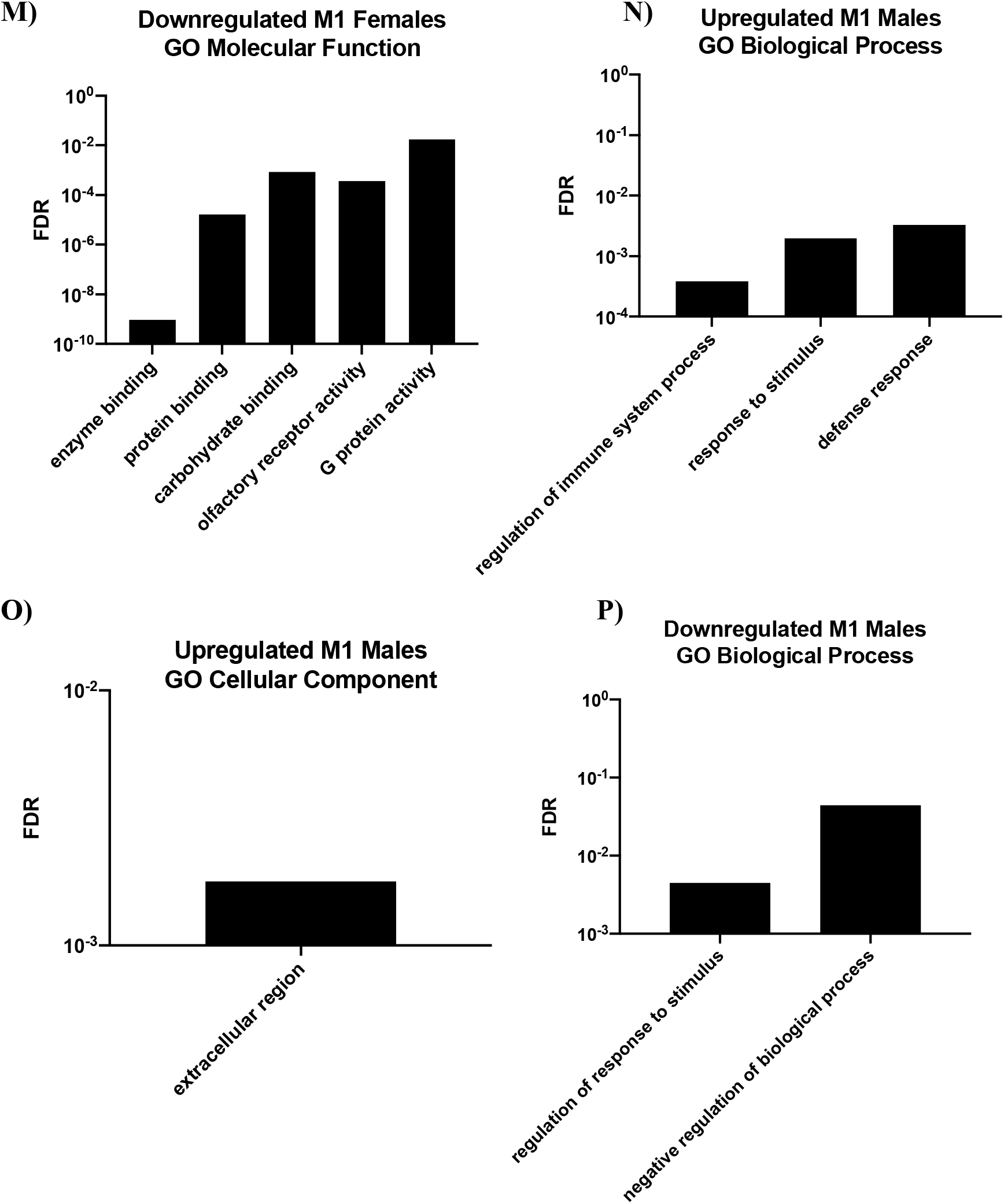
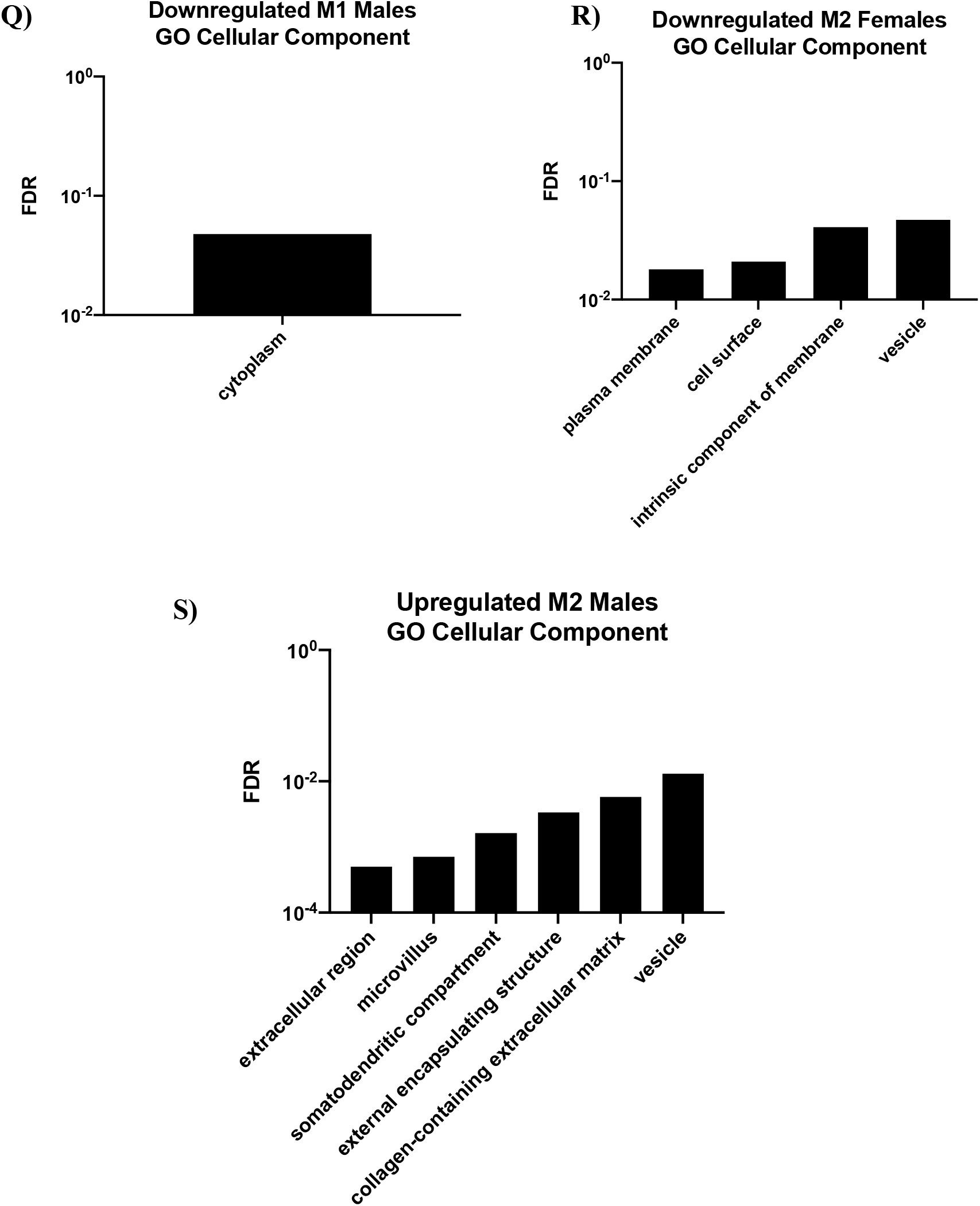
GO Enrichment analysis for up- and downregulated genes induced by iAs exposure for female and male BMDMs across stimulation groups. Plots of GO Enrichment analyses for biological processes, cellular components, and molecular functions were generated for Female M0 groups (A-D), Male M0 groups (E-G), Female M1 groups (H-L), Male M1 groups (M-P), Female M2 groups (Q) and Male M2 groups (R). All enrichment analyses shown on the x-axis are significant (*p*-adj <0.05) and plotted against *p*-adj by FDR correction. If no significant enrichment was found for a comparison group, that group is omitted from this figure. FDR= false discovery rate.

For the upregulated iAs-treated M1 female-derived gene list, enriched biological processes were identified, nearly all of which were related to cell cycle processes (**Fig. 2-6H**); enriched cellular components that followed the same pattern being related to the cell cycle (**Fig. 2-6I**); and enriched molecular functions including protein binding, ATP binding, and cyclin-dependent protein serine/threonine kinase regulator activity (**Fig. 2-6J**). Additional immune-relevant biological processes were identified for the upregulated iAs-treated M1 female-derived gene list, including positive regulation of myeloid cell differentiation, RNA splicing, and cellular response to cytokine stimulus (**Fig. 2-7C**). For the downregulated iAs-treated M1 female-derived gene list, we found enriched biological processes including regulation of localization, inflammatory response, regulation of cell migration and cytokine production (**Fig. 2-6K**); enriched cellular components including cytoplasmic vesicle, endomembrane system, and golgi apparatus (**Fig. 2-6L**); and enriched molecular functions including enzyme binding, carbohydrate binding, and G protein activity (**Fig. 2-6M**). Additional immune-relevant biological processes were identified for the downregulated iAs-treated M1 female-derived gene list, including regulation of vesicle-mediated transport, regulation of phagocytosis, and toll-like receptor signaling pathway (**Fig. 2-7D**). For the upregulated iAs-treated M1 male-derived gene list, we identified significantly enriched biological processes which were regulation of immune system, response to stimulus, and defense response (**Fig. 2-6N**); and one enriched cellular compartment, the extracellular region (**Fig. 2-6O**). No enriched molecular functions or additional enriched biological processes were found for the upregulated iAs-treated M1 male-derived group. For the downregulated iAs-treated M1 male-derived gene list, we identified two enriched biological processes that were regulation of response to stimulus and negative regulation of biological processes (**Fig. 2-6P**); and one enriched cellular compartment, the cytoplasm (**Fig. 2-6Q**). No enriched molecular functions or additional enriched biological processes were found for the downregulated iAs-treated M1 male-derived group.

**Figure 2-7.**
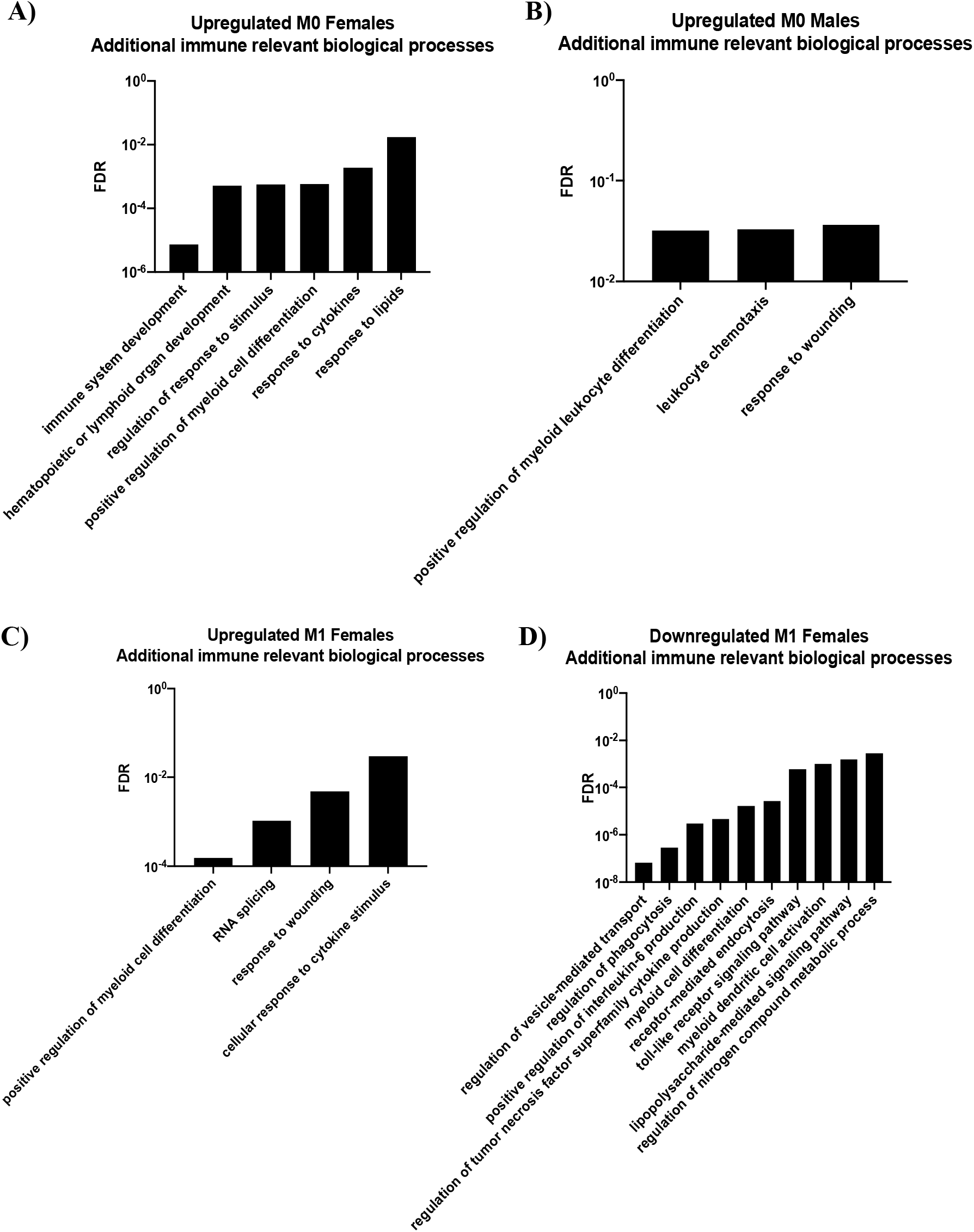
Selected significant immune-relevant GO biological processes for up- and downregulated genes induced by iAs exposure for female and male BMDMs across stimulation groups. Plots of GO Enrichment analyses for biological processes were generated for comparison groups with additional (past the top 10) immune-relevant enriched processes, which included M0 Females (A), M0 Males (B), and M1 Females (C-D). All enrichment analyses shown on the x-axis are significant (*p*-adj <0.05) and plotted against *p*-adj by FDR correction. If no additional immune-relevant significant enrichments were found for a comparison group, that group is omitted from this figure. FDR= false discovery rate.

The upregulated iAs-treated M2 female-derived gene list yielded no significant enrichments for biological processes, cellular compartments, or molecular functions. While the downregulated iAs-treated M2 female-derived gene list showed significant enrichments for cellular compartments, including the plasma membrane and vesicles (**Fig. 2-6R**), no enrichments were identified for biological processes or molecular functions. Like the limited statistically significant enriched findings in the female-derived group, the upregulated iAs-treated M2 male-derived gene list only showed enrichment for cellular compartments including vesicles and collagen-containing extracellular matrix (**Fig. 2-6S**) but no enriched findings for biological processes or molecular functions. For the downregulated iAs-treated M2 male-derived gene list, no enriched findings were found across the three categories.

## DISCUSSION

The unique transcriptomic profiles of iAs-treated BMDMs across stimulation groups, as well as the sex-differences evident within each stimulation group, are clearly demonstrated in our combined RNA-seq and bioinformatics analysis. The raw numbers of perturbed genes (**Figs. 2-2&3**), specifically non-protein-coding genes (**Table 2-15**), transcription factors (**Table 2-16**), and X-linked genes (**Table 2-17**), as well as their likelihood of forming protein-protein interactions (**Table 2-14**; **Fig. 2-3**) and transcriptional regulatory networks (**Fig. 2-3**), suggest that M1-stimulated, especially female-derived, BMDMs are most susceptible to iAs exposure.

M1 macrophages are polarized in response to stimulation by pro-inflammatory cytokines and engagement of PRRs. M1 macrophages are crucial in responding to and orchestrating the inflammatory response against pathogens that cause infectious disease; they release NO to help kill pathogens, are highly phagocytic, and stimulate effector T cells for targeted apoptosis, B cell maturation, and generation of memory (Mills 2000; Orecchioni 2019; Pennock 2013). Thus, iAs-induced immunosuppression of M1 macrophage function may indicate one mechanism by which iAs exposure causes increased risk to infectious diseases such as tuberculosis, lower respiratory tract infections, and influenza (Liao 2011; Rahman 2011; Smith 2011). Our analyses may also help shed light on certain pathways by which arsenic reduces M1 macrophages’ ability to respond to pathogens. The expression of major proinflammatory transcription factors, like IRF8, and its downstream targets were also downregulated in our exposure model in iAs-treated Female M1 BMDMs (**Fig. 2-5C**). Our STRING map categorizations and GO enriched processes are also informative regarding arsenic’s role in perturbing responses to infectious diseases since genes encoding for proteins involved in pattern recognition and antigen presentation, such as TLR7, TLR8, and H2-D1 (Hornung 2008; Watts 2011) had significant PPI scores and were downregulated in STRING network maps generated from the iAs-treated Female M0 and Female M1 downregulated groups (**Figs. 2-4B&E**); on the other hand, inhibitory T cell receptor Ctla4 (Yu 2016) which dampens effector T cell activity and expansion, was upregulated in iAs-treated female-derived M0 and M1 groups (**Figs. 2-4A&D**). Further, GO analysis for biological processes for downregulated DEG lists showed significant enrichments of processes crucial to coordinating a proinflammatory response including leukocyte migration, differentiation, and cytokine and chemokine production and response (**Figs. 2-6&7**). Indeed, we (unpublished data) and others (Kozul 2009) have demonstrated that *in vivo* arsenic exposure causes reduced migration capacity of dendritic cells during influenza infection. Moreover, we observed decreased expansion capacity of macrophages in the lung at 9 days post influenza A virus infection due to arsenic exposure (unpublished data). For the downregulated DEGs in the iAs-treated M1 female-derived group in particular, enrichment was found for genes related to toll-like receptor signaling, LPS-mediated signaling, and regulation of nitrogen compound metabolic process (**Fig. 2-7D**).

Our STRING networks also provide additional general hypothesis-generating mechanisms behind arsenic-induced immunomodulation, as we were able to classify immune-relevant categories across our generated networks which included integrin signaling, coagulation factors, chemotaxis and chemokine signaling, pattern recognition and antigen presentation, splicing machinery, lipid metabolism, complement system, and cytokine signaling (**Fig. 2-4**). These findings are not only relevant in the context of infectious disease (Batista-Gonzalez 2020) but may be implicated in the underlying inflammatory status of many other arsenic-induced diseases like cancer, atherosclerosis, and other cardiovascular diseases where the M1/M2 macrophage polarization dichotomy is well-studied (Batista-Gonzalez 2020; Makhani 2023; Mehla 2019; Lee 2018). For example, macrophages in both the tumor microenvironment and atherosclerotic plaques have been shown to have altered lipid metabolism that promotes disease progression (reviewed in Batista-Gonzalez 2020). Our STRING map generated from the iAs-treated M1 female-derived upregulated gene list show interactions of genes involved in lipid metabolism (**Fig. 2-4D**) and in our identification of significant immune-relevant GO biological processes, the iAs-treated Female M0 upregulated gene set was enriched for response to lipids (**Fig. 2-5A**). Arsenic’s role in altering macrophage lipid metabolism to that of foam cells in atherosclerotic plaques (Lemarie 2008; Lemarie 2015; Negro Silva 2017; Negro Silva 2021) and its role in rewiring macrophage metabolism to become tumor-promoting is being thoroughly investigated by us and others (Cui 2017; Li 2021; Xue 2021). Makhani *et al*, demonstrated that in IFN-LJ-stimulated (but not IL-4 stimulated) bone marrow-derived macrophages, the genes associated with the liver X receptor (LXR) pathway were downregulated by iAs exposure; and subsequent deletion of LXRα from the hematopoietic compartment through bone marrow transplant rescued arsenic-enhanced atherosclerosis in the apoE-/-mouse model (Makhani 2023). However, we did not observe involvement of the LXR pathway in any of the differently activated macrophages upon iAs exposure.

Further, our GO analyses for upregulated genes in iAs-treated M0 male- and female-derived groups show enrichment for biological processes that are implicated in these diseases such as blood coagulation and wound healing (**Figs. 2-6A&E**). The coagulation cascade, for example, is activated during cancer progression to drive angiogenesis, cell migration, and invasion (Hill 2020) and coagulation factors positively correlate with tumor-associated macrophage (TAM) density in the tumor microenvironment (Zhang 2018).

Data by Hung *et al,* suggests that iAs induces the repolarization of classically activated M1 macrophage to M2 macrophages through the autophagy/mitophagy pathway (Hung 2022). However, our findings did not reveal any major differences in the expression of autophagy/mitophagy genes, other than a negative association with Dram2 in iAs-treated M2 female-derived BMDMs (**Table 2-7**).

Another well-documented (Reichard 2010) mechanism of iAs toxicity is its ability to deplete the methyl pool. Methyl metabolism is an important regulator of gene expression, as methyl groups can bind to DNA and histone proteins, typically to repress transcription of downstream targets (Moore 2013). Methyl availability is also tightly linked to the folate cycle and methyl metabolism produces important precursors for the generation of amino acids and adenosine, a precursor of ADP and ATP, and therefore of nucleotides (Ducker 2017). Our Fantom transcriptional networks showed genes involved in methyl metabolism; the upregulated genes in iAs-treated M0 Female network contained *Shmt1*, a serine hydroxymethyltransferase (**Fig. 2-5A**) and the upregulated iAs treated M1 female-derived network contained *Dnmt1*, DNA methyltransferase (**Fig. 2-5B**). Upregulation of these genes could indicate increased methylation of DNA and histones, which generally has a repressive effect on gene expression (Moore 2013), though since arsenic depletes the methyl pool, these results seem contradictory, and it is possible that these changes in methyltransferase expression are compensatory (Reichard 2010). Further, our GO analysis of upregulated genes in iAs-treated M0 male-derived BMDMs was enriched for positive regulation of macromolecule metabolic processes (**Fig. 2-6E**), and the upregulated iAs-treated M1 female-derived gene set was enriched for nucleic acid metabolic processes (**Fig. 2-6H**), with additional enriched molecular functions for ATP binding, nucleotide binding, and nucleoside phosphate binding (**Fig. 2-6J**).

Another noteworthy mechanism of iAs-induced transcriptomic alterations that was evident in our data set was disruption of splicing machinery. The spliceosome is a multi-subunit protein complex that aids in pre-mRNA to mRNA processing by removing introns and facilitating the joining of exons to protein coding regions (Gehring 2021); thus, any changes in the efficiency or efficacy of splicing machinery could result in expression changes of both protein-coding and non-protein-coding transcripts. Our STRING map of the upregulated iAs-treated female-derived M1 gene set showed the protein-protein interactions of several splicing factors (e.g., *Srsfs*) (**Fig. 2-4D**). Fantom transcriptional networks of upregulated genes in iAs-treated M0 females and downregulated genes in iAs-treated M1 females revealed changes in ribonucleoproteins *Hnrnpa1* and *Snrpc*, respectively, which are required for the structural formation of the spliceosome (Chen 2007; Jean-Philippe 2013). Indeed, iAs exposure has previously been associated with changes in splicing patterns especially during carcinogenesis, which contributed to genome instability, epithelial to mesenchymal transition (EMT), and resistance to senescence (Eckstein 2017; Riedman 2019).

Our analysis opens avenues for mechanistic pursual of arsenic-driven sex differences in disease. The efficiency of arsenic metabolism has been shown to differ between males and females, with most studies supporting females as more efficient methylators of arsenic and therefore less sensitive to its effects (Ferrario 2016; Lindberg 2008). Sex differences in arsenic-induced disease risk have been reported in animal (Ferrario 2008; Waalkes 2006) and human studies (Smith 2006; von Ehrenstein 2005) and appear to depend on the disease context. For example, human tuberculosis (a disease already skewed towards males (Rao 2003)) mortality rates upon early-life iAs exposure were higher in males than in females (Smith 2011), whereas *in utero* iAs exposure combined with IAV infection in early-life resulted in additive increases in airway responsiveness in female mice but not in males (Ramsay 2013). Though further statistical analyses need to be performed to measure the strength of association of sex differences in this data set, we show that 24 X-linked genes are dysregulated in iAs-treated female stimulation groups compared to only 3 across the iAs-treated male stimulation groups (**Table 2-17**). The putative biological significance of some of these X-linked genes is supported by our bioinformatics analysis; GATA1 is a characterized transcription factor of hematopoietic cell lineages (Ferreira 2005; Gao 2015), but there is little research concerning its function in macrophages other than regulating the cell cycle and macrophage differentiation (Tanaka 2000). *Gata1* showed up in our STRING map for the upregulated genes in the iAs-treated M0 female-derived group (**Fig. 2-4A**) and in both the STRING (**Fig. 2-4D**) and Fantom (**Fig. 2-5B**) networks generated from the upregulated genes in the iAs-treated M1 female-derived group. In the downregulated gene set from the iAs-treated M1 females, the STRING network showed interactions between X-linked PRRs TLR7 and TLR8 (**Fig. 2-4E**). Fantom further supported the association of these genes in addition to X-linked *Cysltr1*, a leukotriene receptor (Sasaki 2019) (**Fig. 2-5C**).

Overall, this study highlights the sex-specific transcriptomic diversity of macrophages in response to classic polarization stimuli (M1←M0→M2) and low-dose exposure to a common environmental hazard, iAs. Macrophages exist as a diverse population of cells either circulating or resident in virtually all tissues (reviewed in Mass 2023) that function across a broad spectrum of activation stages, far exceeding the polarized (M1←M0→M2) model (reviewed in Guilliams 2015; and Nahrendorf 2017), all with different roles in health and disease. Our results indicate that iAs uniquely affect macrophages in an activation- and sex-specific manner. To further understand the adverse outcome pathways (AOPs) for how exposed macrophages contribute to iAs-induced disease, it will thus be critical to evaluate the impact of iAs on macrophages in a tissue-, activation- and sex-dependent context. This data set provides hypothesis-driven evidence for the targeting of mechanistic pathways involved in iAs toxicity and immunomodulation. Ultimately, such pathway knowledge can be used to generate much needed immunotherapeutic interventions for iAs-induced diseases in exposed populations.

## ACKNOWLEGDEMENTS

This study was funded by National Institutes of Environmental Health Sciences (NIEHS) grants R00ES024808 (F.C.M.S.) and T32ES07141 (E.I.).

## Notes

### Competing Interest Statement

The authors have declared no competing interest.

